# Expansion Microscopy reveals changes in cellular and extracellular structures between healthy and perturbed tendons

**DOI:** 10.64898/2026.04.12.717957

**Authors:** Marcus A. Woodworth, Tristan McDonnell, Xi Jiang, Zizhao Li, Yuna Heo, Mary Kate Evans, Robert L. Mauck, Su Chin Heo, Nathaniel Dyment, Melike Lakadamyali

**Affiliations:** Department of Physiology, Perelman School of Medicine, University of Pennsylvania, Philadelphia, PA, USA; Penn Muscle Institute, Perelman School of Medicine, University of Pennsylvania, Philadelphia, PA, USA; Cellular and Molecular Biology Program, Department of Biology, San Diego State University, San Diego, CA, United States; McKay Orthopaedic Research Laboratory, Department of Orthopaedic Surgery, University of Pennsylvania, Philadelphia, PA, United States; Department of Bioengineering, University of Pennsylvania, Philadelphia, PA, United States; Translational Musculoskeletal Research Center, CMC Veterans Affairs Medical Center, Philadelphia, PA, USA; CReATE Motion Center, CMC Veterans Affairs Medical Center, Philadelphia, PA, USA; Department of Cell and Developmental Biology, Perelman School of Medicine, University of Pennsylvania, Philadelphia, PA, USA; Epigenetics Institute, Perelman School of Medicine, University of Pennsylvania, Philadelphia, PA, USA

## Abstract

Tendons transmit mechanical forces between muscles and bones through their highly aligned, collagen-rich extracellular matrix. When damaged, resident cells help restore this matrix. However, in tendinopathies, this repair response fails, leading to loss of proper tendon function. How altered mechanical states reshape tendon cell and matrix architecture remains poorly understood because existing methods do not readily capture tendon structure across relevant length scales. Here, we combine Fluorescent Labeling of Abundant Reactive Entities (FLARE) with Expansion Microscopy (ExM) to visualize cellular and extracellular structures in native tendon tissue. FLARE-ExM resolves the dense fibrillar matrix across multiple tendon types, including elastic fibers and glycan-rich cellular protrusions. In a tendon resection model, acute loss-of-tension was associated with increased fibril width and expansion of carbohydrate- and protein-rich regions. In ruptured human Achilles tendon, FLARE-ExM revealed extracellular disorganization and disrupted cellular architecture. These results establish FLARE-ExM as a useful approach for studying how mechanical perturbation remodels tendon architecture across physiological and disease contexts.

## Introduction

Tendons transfer tensile mechanical forces between muscles and bones. Proper transmission of these forces requires a highly organized extracellular matrix (ECM) composed mostly of tightly packed, parallel collagen fibrils that come together to form larger fiber bundles (*1*). Cells within this tissue, termed tenocytes, secrete ECM components and catabolic regulators that maintain and restore tendon tissue structure (*2*). Given their location, these cells are subject to numerous biomechanical and physiochemical signals (*3*) through thousands of daily loading cycles generated by muscle contractions (*4*). However, tendon overuse can cause tissue degeneration (e.g., tendinopathy), where local microdamage disrupts ECM integrity. Cells respond to this microtrauma by producing proteoglycans and matrix remodelers that can disorder the collagen matrix (*5–7*), leading to a local loss of mechanical cues (*8*, *9*). Common treatment for these cases involves physical therapy (*10*), aimed at stimulating resident cells with controlled mechanical cues to promote tissue regeneration. However, about a third of those treated will not fully recover (*11*) since loss of mechanical cues causes cells to develop aberrant behaviors, such as chondrogenesis (*12*), exacerbating pathogenesis. Understanding how altered mechanical cues reshape tendon cell and matrix architecture is therefore important for explaining loss of cell function and for developing more effective therapies.

Optical microscopy has enabled visualization of tendon structures across multiple length scales. For example, confocal microscopy is a high-volume imaging technique (*13*), which, when combined with second harmonic generation (SHG) and 2-photon microscopy, has enabled visualization of collagen and elastic fiber organization throughout tendons (*14–16*). Using these imaging techniques, researchers have established a hierarchical organization for tendons in larger animals (e.g., human and horse), where collagen fiber bundles form fascicles encapsulated by an interfascicular matrix (IFM) (*17*), composed of cellular and extracellular structures that differ from the fascicular matrix (FM) (*18*). Rodents, model organisms commonly used in tendon physiology and pathology studies, lack distinctive IFM and FM regions, presenting an architecture composed of interconnected cells that encapsulate nearby fiber bundles (*19*). Together, these studies define important aspects of tendon organization, but they do not provide nanoscale, molecularly informative imaging of dense tendon architecture in intact tissue.

Electron Microscopy has been instrumental in determining the nanoscale structures of tendons. For example, Scanning Electron Microscopy (SEM) provides details on the morphology of collagen fibrils at the nanoscale level, while Transmission Electron Microscopy (TEM) can reveal fibril width and packing (*20*). These methods, however, have limited depth capabilities and are normally applied only to thin sections or surface view of fibrils only. To obtain further 3D details of the ECM, previous work has employed focused ion beam and scanning electron microscope (FIB-SEM) or Serial Block Face on tendons, allowing the segmentation and tracking of fibrils throughout the tissue (*21–24*). These techniques have allowed the visualization of tenocytes and fiber organization at various stages of tendon development (*25*). However, EM relies primarily on contrast alone to identify structures, leaving much of the non-collagenous ECM, such as elastic fibers and proteoglycans, hidden. Moreover, the labor-intensive nature of sample preparation and imaging inherently limits throughput, limiting both the number and total area of tendons that can be visualized in each experiment. Thus, a major unmet need is a method that can resolve tendon ultrastructure in 3D while retaining molecular contrast and sufficient throughput for comparing physiological and pathological conditions.

Expansion microscopy (ExM) enables multiplexed labeling and visualization of structures in thick tissues at resolutions beyond the diffraction limit of light (*26*). ExM anchors tissue structures to a swellable gel which, after disruption of rigid structures, allows for isotropic expansion of the sample in water, achieving super-resolution by physical magnification. ExM works with conventional antibodies (*27*, *28*), *in situ* labeling of nucleic acids (*29*), and other various cell and tissue samples (*30*, *31*), converting a conventional fluorescence microscope into a super-resolution one without the use of specialized equipment or dyes. A recent extension of ExM, here referred to as Fluorescent Labeling of Abundant Reactive Entities (FLARE), uses covalent labeling of abundant functional groups to generate bright, feature-rich images of tissue architecture without antibodies (*32*, *33*).

Although most ExM techniques have been predominantly applied to either cultured cell lines or softer tissues (e.g., brain), various attempts have been made to adapt the protocol to denser tissue types. For example, ExM has been employed to expand collagen-rich tissue sections of the skin and the cornea (*34*), whole zebra fish and mouse embryos (*35*), and most recently even a whole neonatal mouse has been expanded (*36*). However, ExM has not, to our knowledge, been applied to resolve nanoscale cellular and extracellular architecture in dense mature tendon, particularly in older mice and human fibrous connective tissue. This is likely because musculoskeletal tissue is stiffer than most other tissues, particularly in older organisms (*37–39*), which can hinder proper expansion. Since various musculoskeletal pathologies manifest later in life, there is a great need to adapt super-resolution microscopy techniques, such as ExM, to more mature and dense tissue types, including human fibrous connective tissues.

To address this gap, we established a tendon tissue optimized FLARE-ExM workflow that enables nanoscale visualization of cellular and extracellular architecture in native tendon tissues. Across multiple tendon types and species, FLARE-ExM resolved collagen fibrils together with non-collagenous structures, including elastic fibers and glycan-rich cellular protrusions. In mouse Achilles tendon, it revealed a complex elastic fiber network throughout collagen fiber bundles and the sheet-like protrusions of nearby cells. These protrusions, in turn, presented glycoprotein-rich connections between neighboring cells, as they wrapped around nearby fibers. In a *Flexor Digitorum Longus* tendon resection model (*40*), loss of tension was associated with increased fibril width and expansion of carbohydrate- and protein-rich regions. Lastly, we applied FLARE-ExM to ruptured human Achilles tendon, revealing disordered ECM throughout the tissue, hypervascularization, and chromatin enrichment at the nuclear periphery. Together, these results position FLARE-ExM as a powerful approach for studying how mechanical perturbation remodels tendon architecture across scales.

## Results

### Adapting FLARE-ExM to study dense tendon tissue ultrastructure

To enable expansion microscopy of dense mature tendon, we optimized FLARE-ExM for intact tendon explants and cryosections (Fig. 1). We found that although the original FLARE-ExM protocol, which employs the MAP gel solution to link proteins to the gel and denaturation solutions to passivate the sample (*41*), expands 50-µm cryosections of the bovine Achilles tendon, the protocol had difficulty in expanding mouse tail tendon explants. These tendons either failed to expand in the direction parallel to the longitudinal axes of the tendon (Fig. S1A) or would detach from the gel before full expansion, possibly due to their stiff composition. We therefore modified the protocol to control the sample expansion rate during and after sample denaturation. Following the protocol used for C. *elegans* expansion, which reported similar detachment problems (*42*), we added higher salt concentrations as well as calcium ions to the denaturation buffer to help mitigate expansion during denaturation. In addition, we used step-wise dilution of salts during the final expansion step, enabling a slower expansion rate for the sample, as previously done for the expansion of collagen-rich skin or corneal tissues (*34*). These adaptations, along with the use of thinner gels to facilitate solution exchange, allowed for preservation of the tendon explant within the gel during expansion, and demonstrated a uniform expansion in mouse tail tendons (Fig. S1B). Minor- and major-axis nuclear expansion factors of 4.5× and 4.7× were similar to each other and to the bulk gel expansion factor (4.6×), consistent with near-isotropic expansion (Fig. S1C).

**Fig. 1.**
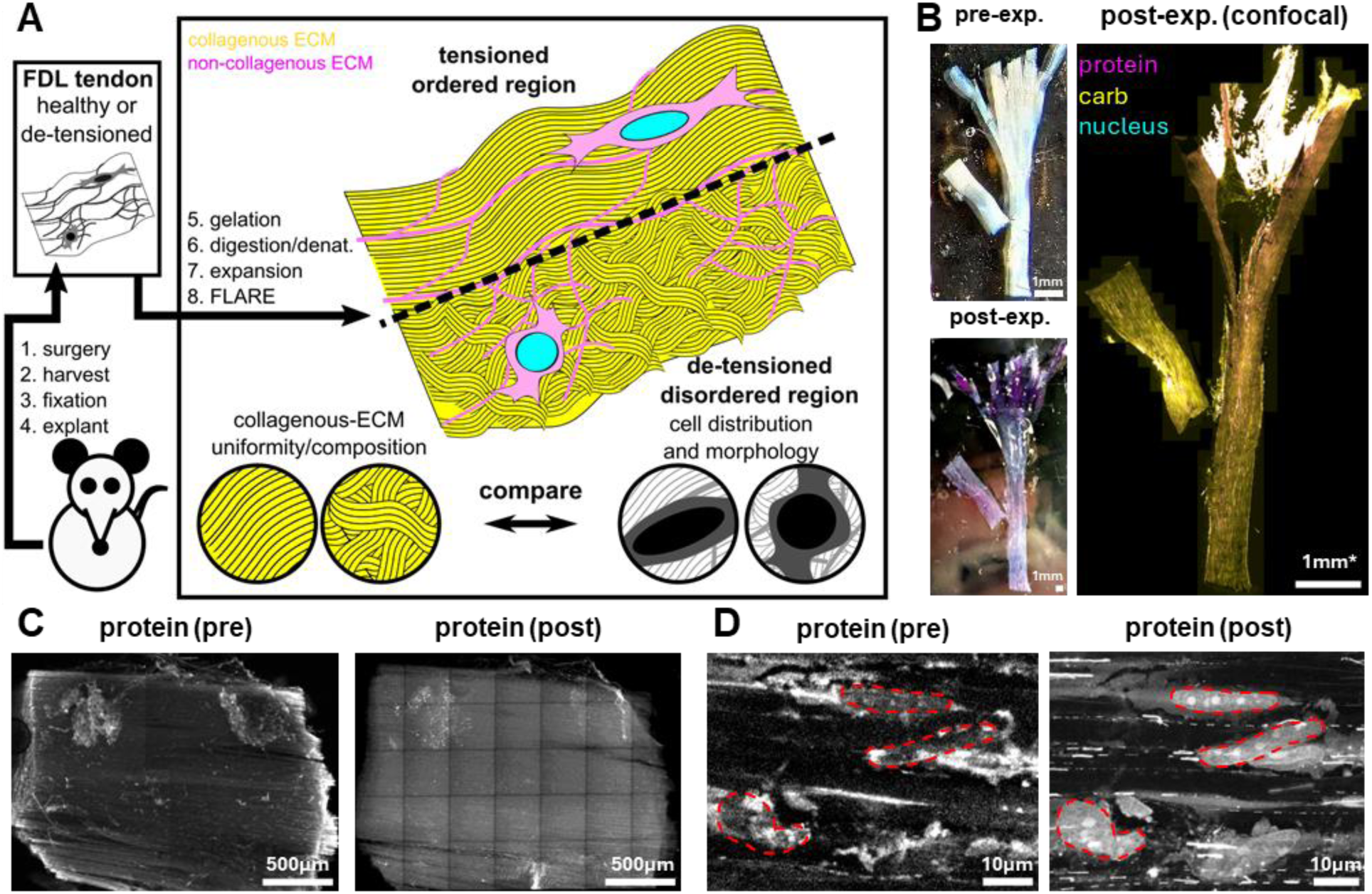
FLARE-ExM of tendon tissue workflow. **A.** Example workflow for a mouse tendon explant treatment and expansion. A mouse *Flexor Digitorum Longus* (FDL) tendon surgically resected. The mouse limb is later harvested and fixed over several days in formalin, and the FDL tendon is excised. The tendon explant is then incubated in monomer for a few days, digested and/or denatured over several days, treated with FLARE and then expanded for ultrastructure analysis of tendon uniformity and composition. **B.** Example images of an FDL tendon explant after gelation (pre-expansion), after FLARE and expansion (post-expansion), after imaging on a confocal (post). Scale bar for the confocal (post) image has been corrected to pre-expansion units. **C.** Correlative imaging of a whole mouse patellar tendon explant FLARE-stained for proteins before expansion (left) and after expansion (right), maximum intensity projected over 120 µm. **D.** High-magnification image, maximum intensity projected over 4µm of cells within the pre- and post-expanded patellar tendon explant in **(C.)**, with a red outline indicating the same traced cell nuclei perimeter between pre- and post-expansion. Images in (C.) were subject to a 2-pixel radius gaussian filter.

Applying this optimized protocol to other mouse tendon explants, including the Achilles, plantaris, patellar, and *flexor digitorum longus* (FDL) tendons (Table 1), we confirmed that all tendon types remained stably embedded in the gel following expansion (Fig. 1B). Notably for larger and longer tendons (i.e., Achilles and FDL), we noticed sections of “crinkling” within the tendon, particularly in thicker tissue areas (Fig. S1D). Although these artifacts resembled those reported after fixation or freeze-thaw cycles (*20*) and may not be due to the expansion itself, we therefore incorporated a collagenase type VII digestion before denaturation, a step previously employed by others (*35*, *42*), along with an additional elastase digestion after denaturation, to further passivate the tissue and minimize any potential structural distortions. The digested tendons expanded comparably to undigested samples, and although the FLARE signal intensity decreased within the ECM region, the tissue exhibited a more homogenous appearance with reduced “crinkling” overall.

**Table 1.** Differences in digestion and denaturing conditions for expanded tendon samples.

| **Tendon sample** | **Denaturation solution** | **2<sup>nd</sup> denaturation step time (days)** | **Enzyme digestions** |
| --- | --- | --- | --- |
| Achilles (Bovine) | MAP | 3 | - |
| Tail | ExCel | 1 | - |
| Patellar | ExCel | 3 | - |
| Plantaris | ExCel | 3 | - |
| FDL | ExCel | 3 | - |
| FDL (digested) | ExCel | 3 | Coll. (VII),* Ela. (II) † |
| Achilles | ExCel | 3 | - |
| Achilles (digested) | ExCel | 3 | Coll. (VII),* Ela. (II)† |
| Achilles (Human) | ExCel | 3 | Coll. (VII), * Ela. (II),†<br>chon. ABC‡ |

\*Coll. (VII): Collagenase type VII, †Ela. (II): Elastase type II, ‡chon. ABC: Chondroitinase ABC

We further evaluated the expansion uniformity using correlative imaging between pre- and post-expanded tendons, with and without the digestion steps (Fig. 1C and Fig. S2). Comparison of the same fields of view before and after tendon expansion showed high correspondence of protein structures, with a brighter protein FLARE signal achievable in the non-digested sample. Accordingly, we applied these protocol adaptations as appropriate, based on tissue source and thickness—for example, using the MAP denaturation solution for bovine tendon cryo-sections, the calcium-containing denaturation solution for tendon explants, and incorporating digestion steps for thicker tendons such as those for the Achilles and FDL (Table 1).

### FLARE-ExM highlights the densely packed and parallel collagen fibrils of tendon cryosections

To assess whether FLARE-ExM resolves key tendon ECM features, we first examined bovine Achilles tendon cryosections as they expanded with ease using the original, unmodified MAP protocol solutions with longer incubation times (Fig. 2A). The protein and nuclear stains of FLARE highlighted the cytoplasm and the nucleus of resident tendon cells, the majority of which are tenocytes, while the carbohydrate stain resolved a dense, parallel fibrillar network consistent with tendon collagen organization. To accurately measure the width of these fibril-like structures and directly compare our values with those reported from TEM analyses, we rotated a section of the gel 90° to image the sample in a horizontal view (Fig. 2B, top). In this view, the fibril-like structures we observed appeared as densely packed puncta (Fig. 2B, middle) with mean widths of 118 ± 82 nm and 116 ± 81 nm for x and y, respectively, consistent with isotropic expansion (Fig. 2B, right). This width closely matched values previously determined for collagen fibrils in mouse tendons (*21*) and Bovine Tendons (*43*). We speculate carbohydrate signal likely corresponds to the proteoglycans and glycosaminoglycan (GAGs) present on collagen fibrils (*44*), which allows us to the visualize the tendon fibrillar network.

**Fig. 2.**
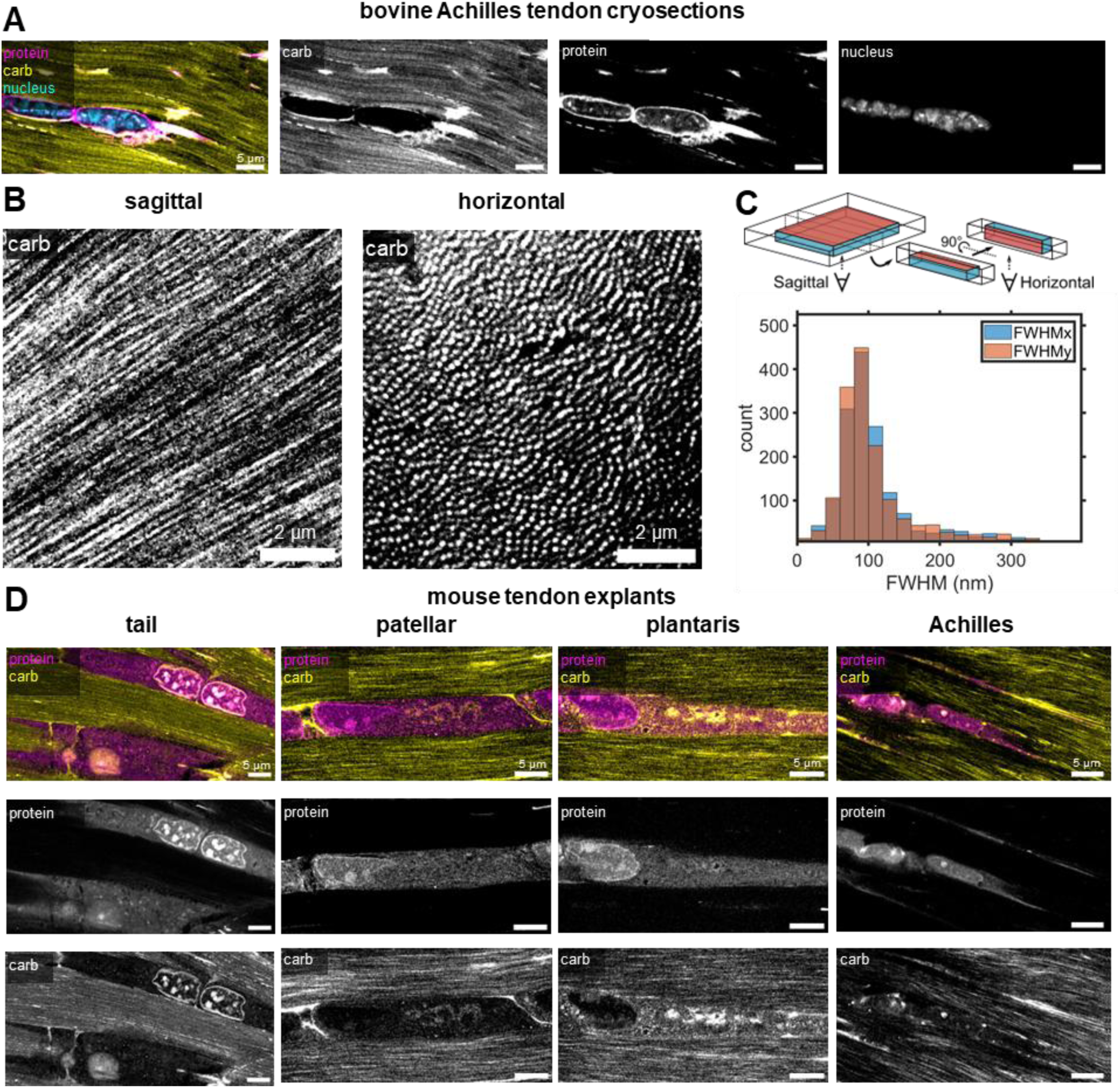
FLARE reveals meso- and nano-scale features of the extracellular matrix across different tendon tissue types and species. **A.** Sagittal view of a 50 µm cryosection expanded bovine Achilles tendon with FLARE-stained proteins (magenta), carbohydrates (yellow) and nuclei (cyan). **B.** Sagittal **(left)** and horizontal **(right)** views of the carbohydrate signal in a FLARE-stained 50 µm cryosection bovine Achilles after an extended (4hr) oxidation step. **C. Top.** Diagram for how an expanded tendon can be imaged in a sagittal view, then after slicing a section of the gel and rotating it 90°, can be imaged in a horizontal view. **Bottom.** Histogram of the x and y Full Width Half Max (FWHM) of the 2D gaussian fit for the detected puncti of the tendon imaged in the horizontal view across a larger field of view than shown in **(B.). D.** Example sagittal views for four different expanded mouse tendon tissues with FLARE-stained proteins (magenta) and carbohydrates (yellow) at the cell level. Images in **(B.)** and **(D.)**, were subject to a 2-pixel radius gaussian filter.

In addition to the carbohydrate-rich fibrils seen throughout the ECM region, we identified a subset of fibers that were distinctly enriched in both protein and carbohydrate components, presenting a dashed-line appearance (Fig. S3). Expansion is likely responsible for the dashed appearance of these protein and carbohydrate enriched structures, as they appear more continuously in the unexpanded samples (Fig. 1C and Fig. S2). Given their proximity to cells but difference in signal compared to the dense collagen fibril network, we hypothesized these structures corresponded to elastic fibers (*45*). These fibers, which comprise up to 10% of the tendon (*46*), are suggested to play a role in the stability of the interfascicular matrix (*47*), and the mechanical function of the fascicular matrix (*16*). Elastin immunolabeling indeed showed strong spatial overlap with these protein- and carbohydrate-enriched fibers (Fig. S3). These results demonstrate that FLARE-ExM can reveal specific non-collagenous features of the ECM, without the need for antibody-based labeling.

### FLARE-ExM reveals the ultrastructure of various mouse tendon explants and adjacent tissues

We next applied FLARE-ExM to a variety of mouse tendons, adopting the protocol as needed (Table 1) to examine the extracellular and cellular features in more detail (Fig. 2D, Fig. S4A). All tendon types showed similar tenocyte organization and fibrillar ECM architecture. In addition, quantification of fibril widths within the ECM region revealed a high degree of consistency across the different tendon types and expansion protocols (Fig. S4B). Together, these data show that FLARE-ExM can resolve tendon ultrastructure across multiple tendon types and species.

Since explanted tendons were obtained from fixed mouse-limbs, we could visualize additional tissue types neighboring each tendon (Fig. S5). Of note, we visualized the sarcomeres of muscles (Fig. S5A), along with an enrichment in carbohydrate signal at the myotendinous junction (Fig. S5B), likely due to the high abundance of sarcoglycans, dystroglycans and laminins at this region (*48*). The carbohydrate stain also labeled a large vessel adjacent to the Achilles tendon, showing invaginations of the vascular smooth muscle nuclei (Fig. S5C&D). These results demonstrate that FLARE-ExM not only resolves the fine architecture of tendon-specific regions but also visualizes structural interfaces between tendon and neighboring musculoskeletal tissues.

### Tendon ECM contains a network of elastic fibers that link to interconnected cell sheets

Since the Achilles tendon is a clinically relevant target, as it has a high prevalence of developing tendinopathies in humans (*49*, *50*), we decided to visualize the 3D organization of this tendon in greater detail (Fig. 3). We applied FLARE-ExM, with collagenase and elastase digestion, to a mouse Achilles tendon explant. In the horizontal view, protein-rich tenocyte cell bodies formed sheets that connected neighboring cells and wrapped around collagen fiber bundles (Fig. 3A). This cellular organization closely resembled that seen previously by SBF-SEM, in which tenocytes interconnect with neighboring cells early in development, and form sheets that wrap around adjacent fiber bundles as the tissue matures (*25*). Interestingly, we found patches of high carbohydrate intensity within these regions, particularly between cells at the cell and fiber boundaries (Fig. S4A). These regions may correspond to glycoproteins involved in tendon cell adhesion, such as adhesion-promoting proteoglycans (*51*) , and cell-cell junctions, such as cadherins (*52*, *53*).

**Fig. 3.**
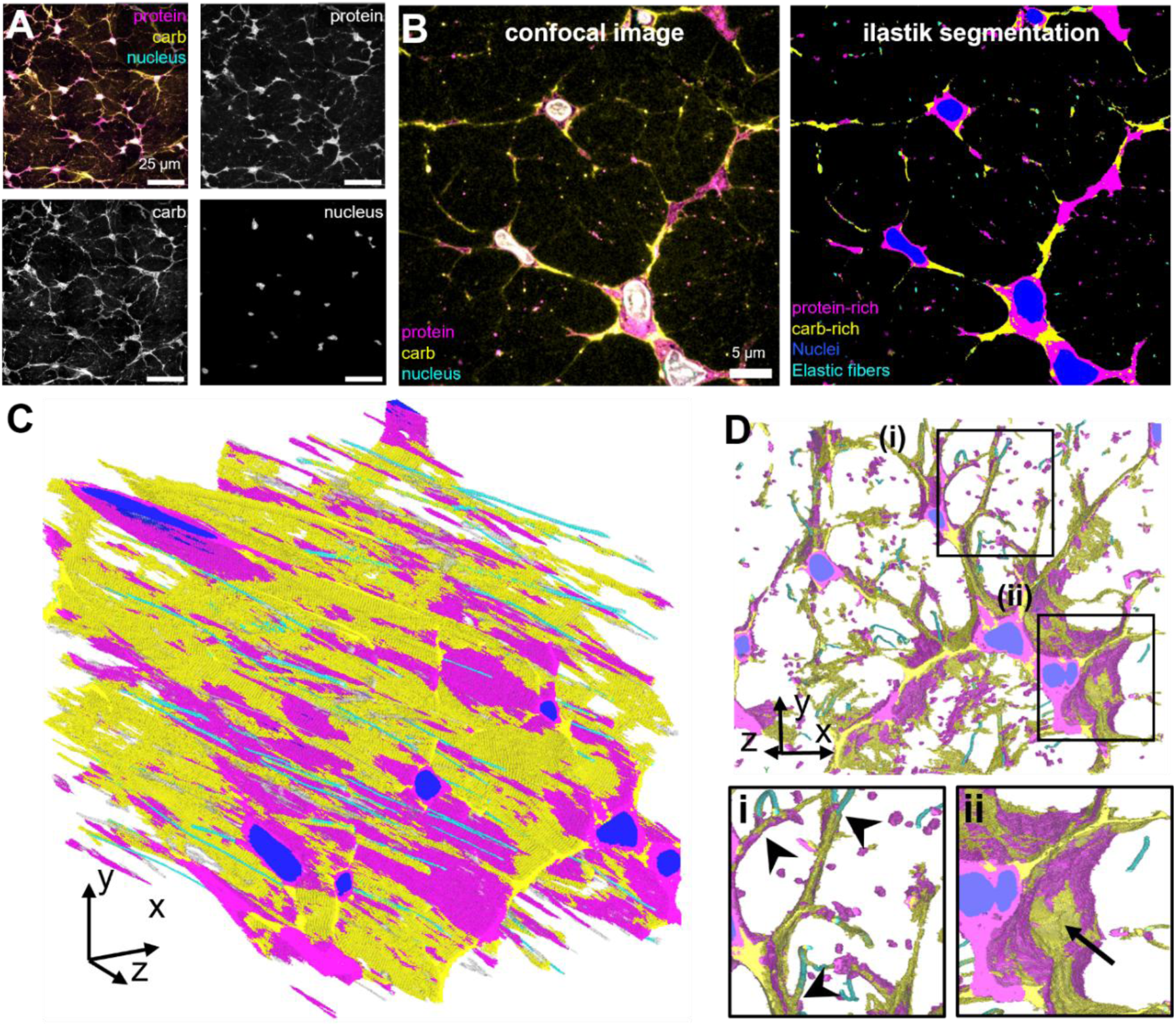
Tendon fascicles have protein-rich and carbohydrate-rich sheets that encompass tendon fibril bundles. **A.** Horizontal view of an expanded Mouse Achilles tendon explant, with FLARE-stained carbohydrates (yellow), proteins (magenta), and nuclei (cyan). **B.** Example slice of a 3D confocal stack of the expanded tendon in (A.) on the left, with the ilastik segmented nuclei (blue), protein-rich regions (magenta), carb-rich regions (yellow) and elastic fibers (cyan). **C.** 3D image of the ilastik segmented regions of the FLARE-ExM treated tendon in **(A.)** showing the spatial distribution of segmented nuclei (blue), protein-rich regions (magenta), carb-rich regions (yellow) and elastic fibers (cyan). **D.** 3D image from **(C.)** oriented to see down the fiber bundles, along with zoomed in views of two regions showing elastic fiber connections to the protein-rich/carbohydrate-rich regions (arrowheads) in **i.**, and a carbohydrate-rich patch (arrow) of the cellular region in **ii.** Images in **(A.)** and **(B.)**, were subject to a 2-pixel radius gaussian filter.

Interspersed throughout both the cellular and fiber regions of the Achilles tendon were puncta of high carbohydrate and protein signals, with a subset of these regions appearing as ringed structures (Fig. S4B). Based on their FLARE signature and elastin immunolabeling (Fig. S3), as well as the results seen by others (*15*, *45*), these puncta likely represent elastic fibers. To understand how these regions were organized in 3D space, we obtained a stack of the tendon in the horizontal view and used ilastik (*54*) to segment the carbohydrate- and protein-rich regions, and elastic fibers present within the different fiber bundles (Fig. 3B&C, Movie S1). This analysis revealed that the carbohydrate-rich patches seen in 2D span across the length of the cell matrix, particularly between protein-rich regions surrounding cell nuclei (Fig. 3Cii) and at the border of the collagen fiber bundles and cells. In this view, we also found that elastic fibers branched at different points (Fig. S5C) and connected to the sheet-like protrusions of cells (Fig. 3Ci), showing deep extensions at times into these sheets (Fig. S5D). FLARE-ExM therefore reveals an elastic-fiber network embedded within collagen fiber bundles, which are in turn surrounded by the tenocyte matrix proximal to each bundle.

### Loss-of-tension results in increased fibril widths and carbohydrate- and protein-rich regions

To test how mechanical cues change the cellular and extracellular architecture of a tendon, we employed a *flexor digitorum longus* (FDL) tendon resection model (*40*). In this model, the FDL tendon is resected in the mouse forefoot to acutely de-tension the tendon in the proximal region, allowing us to examine how acute loss of tension alters tendon architecture proximal to the tibia while minimizing contributions from early wound healing at the resection site. For our study, we retrieved FDL tendons from 1- and 6-day post-surgery mice from the uninjured hind-limb, and surgically resected hind-limb. Each pair of FDL tendons were processed side-by-side within the same gel to help maintain the same expansion and staining conditions per mouse. Comparing these two tendons for each mouse, we observed a modest increase in fibril width for Day 1 tendons and a more pronounced increase (from 116 ± 25 nm to 137 ± 30 nm) for Day 6 tendons (Fig. 4C). We also detected enrichment of carbohydrate stain for the resected tendon compared to the uninjured control (Fig. 4D, sup. Fig. 7&8). Horizontal views further demonstrated increased nuclear cross-sectional areas (Fig. 4E), without changes in nuclear circularity (Fig. S10A), along with an increase in carbohydrate and protein-rich area fraction in and around cells (Fig. S10B), which likely correspond to larger cell cytoplasmic and membrane-associated glycoprotein regions. Despite these changes, we did not detect an increase in branch points proximal to the nuclei in resected tendons (Fig. 4F).

**Fig. 4.**
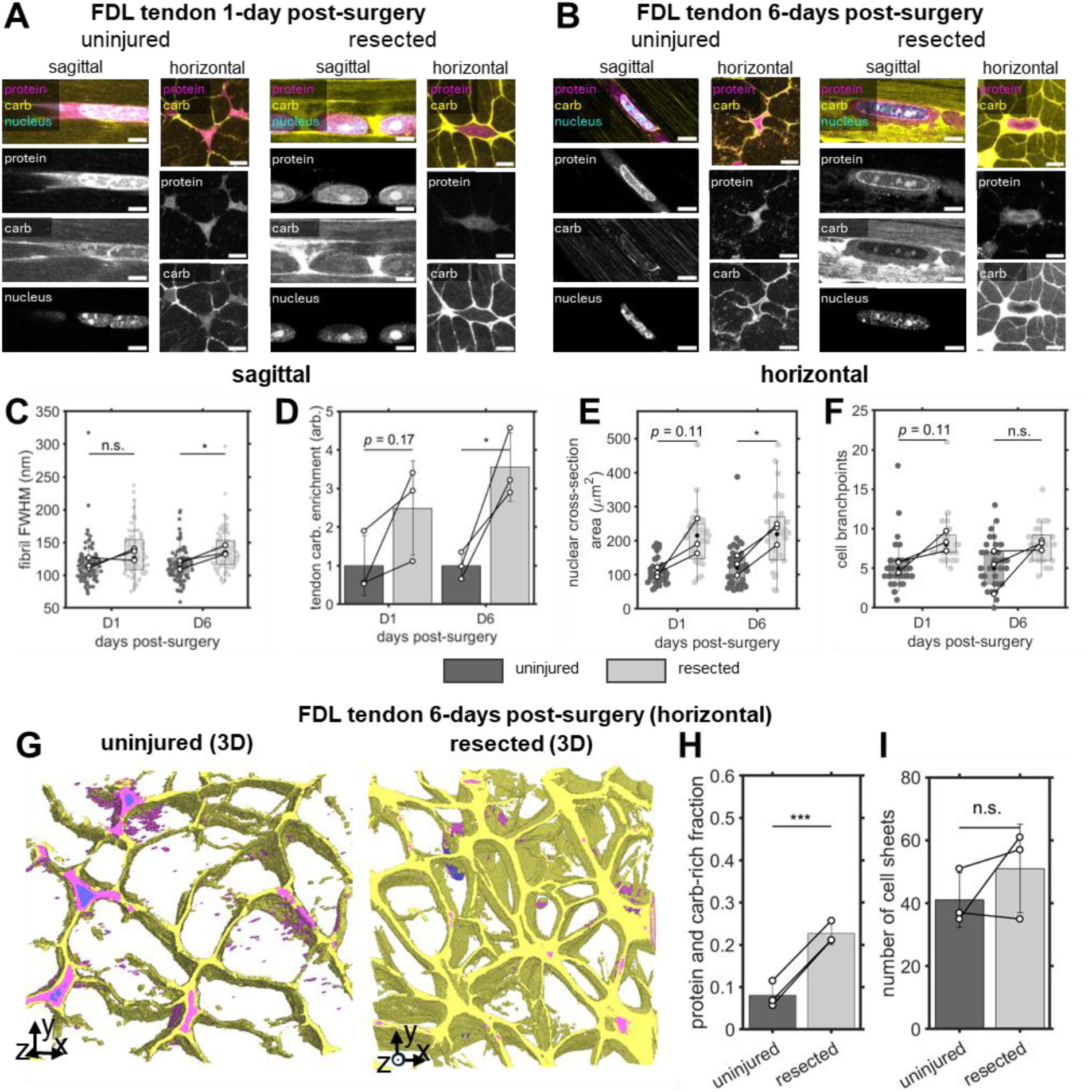
De-tensioned *flexor digitorum longus* (FDL) tendons show larger carbohydrate and protein sheet area compared to uninjured tendons. A. and. **B.** Sagittal and horizontal views of expanded tenocytes within uninjured (left) and resected (right) FDL tendon explants 1 day **(A.)** or 6 days **(B.)** post-surgery, with FLARE-stained carbohydrates (yellow), proteins (magenta), and nuclei (cyan). **C.** Full width half max (FWHM) of carbohydrate-stained fibrils of either uninjured or resected FDL tendon explants 1 day (D1) or 6 days (D6) post-surgery. **D.** Carbohydrate enrichment within the collagenous ECM region of uninjured or resected FDL tendon explants 1 day (D1) or 6 days (D6) post-surgery. **E.** Nuclear area determined for cells from uninjured or FDL tendon explants D1 or D6 post-surgery, imaged in a horizontal view. **F.** Number of branches from a skeletonized image of the cellular region of either uninjured or resected FDL tendon explants D1 or D6 post-surgery, imaged in a horizontal view. **G.** Example of FLARE-stained 3D image stacks of uninjured (left) or resected (right) FDL tendon explants 6 days post-surgery, with ilastik segmented nuclei (blue), protein-rich (magenta) and carb-rich (yellow) regions. **H.** Fraction of the protein- and carb-rich regions, which mostly correspond to cellular volume, for the uninjured or resected FDL tendon 6 days post-surgery **I.** The number of cell sheets observed within the imaged tendon volume in uninjured or resected 6 days post-surgery FDL tendon explants. Sagittal images in **(A.)** and **(B.)** were subject to a 2-pixel radius gaussian filter, while the horizontal images were subject to a 2-pixel radius median filter.

Given the variability in tendon carbohydrate and protein-rich area fraction for the day-6 post-surgery tendons (Fig. S10B), we decided to obtain volumetric data for these tendons in a horizontal view (Fig. 4G, Movie S2). In doing so, we confirmed that the carbohydrate and protein-rich volume fraction increased in 3D space (Figure 4H), but without changing the number of sheets within the field of view (Figure 4I). These results suggest that acute loss of tension is accompanied by expansion of glycoprotein-rich regions within fiber bundles, without an obvious change in the number of cell-sheet connections in the sampled volume.

### FLARE-ExM highlights cellular and extracellular structural abnormalities in human ruptured Achilles tendon

Although histological images have been obtained for ruptured human Achilles tendons (*55*), the nanoscale cellular and extracellular organization remains poorly characterized. We therefore sought to apply FLARE-ExM to 100 µm cryosections of a ruptured human Achilles tendon sample. Human ruptured tendon sections were more resistant to uniform expansion. We therefore added chondroitinase ABC during the elastase digestion step to mildly digest proteoglycans, given their implication in crosslinking collagen fibrils (*56*). This small modification resulted in a much more homogenous staining pattern in the tendon section (Fig. 5A). We observed extensive hypervascularization and hyper-cellularization, consistent with prior histological descriptions of ruptured Achilles tendon (*55*). Visualizing the tendon at the cellular level, we saw extensive disorder of the collagen fibers between cells (Fig. 5Bi), along with a lack of a cohesive, interconnected cell network (Fig. 5Bii). Additionally, the cell nuclei were no longer elongated, but rather showed either a rounded morphology, or multiple irregular protrusions (Fig. 5Biii). Upon inspection of a single nucleus at different z-planes (Fig. 5C), we observed these protrusions were connected to a single nucleus. In addition, there was high enrichment of protein signal, likely corresponding to chromatin, at the periphery of nuclei, distinct from the more uniform distribution observed in mouse and bovine Achilles tendon nuclei and closely resembling the chromatin organization of isolated cells from tendinosis patients or from tenocytes grown on soft substrates (*57*). This data, to our knowledge, provides the first nanoscale views of cellular and nuclear architecture in ruptured human Achilles tendon.

**Fig. 5.**
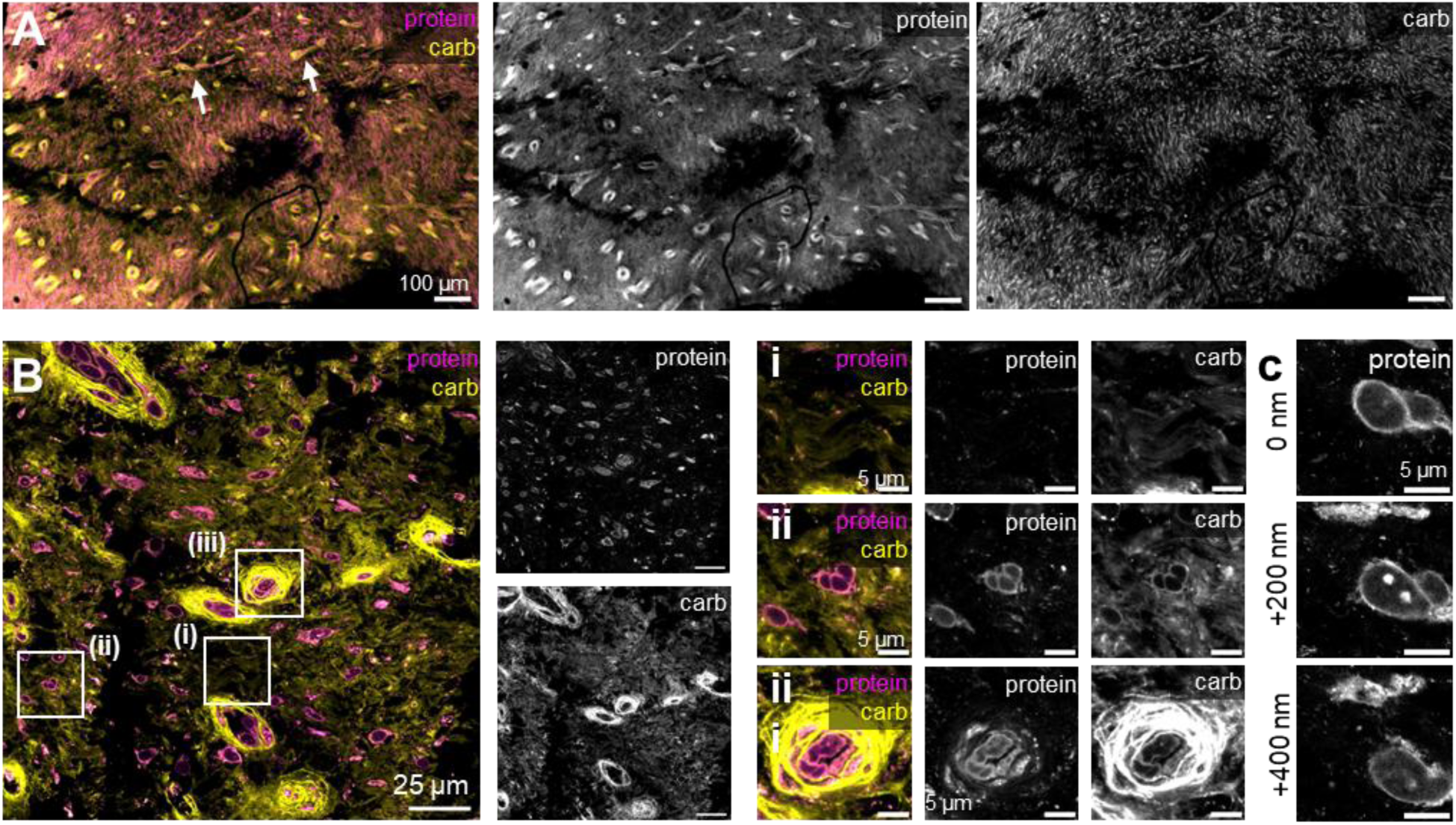
FLARE on Ruptured human Achilles tendon cryosections reveals hypervascularization, ECM disruption, and changes in nuclear morphology. **A.** FLARE staining of protein (magenta) and carbohydrates (yellow) of a ruptured human Achille tendon 100 µm cryosection, with arrows indicating regions of hypervascularity. **B.** High-magnification tile scan of the tendon sample in **(A.)** with zoomed in views showing ECM disruption in **i.,** an irregular and multi-lobular shaped nucleus in **ii.**, and microvasculature in **iii. C.** (top) FLARE-stained protein of a tenocyte cell imaged at three different depths, showing irregular nuclear morphology and protein signal enrichment at the nuclear periphery. All images were subject to a 2-pixel radius gaussian filter.

## Discussion

We adapted the super-resolution technique, ExM, together with a broad labeling strategy, FLARE, to interrogate architectural components within intact tendon tissue explants at resolutions beyond the diffraction limit of light. This methodology is highly robust, being applicable to tissue sections and explants from several types of tendons across several species. Historically, visualization of nanoscale tendon structures such as collagen fibers has relied primarily on electron microscopy, an approach limited by low throughput, low contrast, and low molecular specificity, making it challenging to study how tendon architecture changes during disease progression. FLARE-ExM enables visualization of not only the fine structures of collagen fibrils for multiple tissue types, but also of non-collagenous features, including elastic fibers and their spatial relationship to interconnected cellular regions that wrap around fiber bundles. Using FLARE-ExM, we demonstrated that loss of mechanical tension is associated with increased glycoprotein accumulation throughout the tendon, thickening of collagen fibrils, and an increase in carbohydrate-rich regions over whole tendon space.

Changes in collagen fibril width are a hallmark of both tendon development and a range of tendon pathologies. Quantitative measurements of fibril width have traditionally relied on TEM or SBF/FIB-SEM, techniques that are labor-intensive, low-throughput and less accessible than most light-microscopy-based methods. In this study, we demonstrate that FLARE-ExM can be used to quantify collagen fiber widths within intact tendons using a standard confocal microscope. For sufficiently small tendons, this approach did not require physical sectioning prior to gelation and expansion to obtain sagittal and horizontal views of the tendon architecture. Instead, horizontal views could be obtained by coarse sectioning of the same expanded gel initially imaged in the sagittal orientation. This flexibility highlights the practical advantages and experimental versatility of working with expanded samples, allowing for imaging of tendon ultrastructure across the entire length of a tendon, and as small as the width of individual fibrils.

It is important to note that the structures detected by carbohydrate staining within the collagenous-ECM likely reflect proteoglycans or GAGs associated with collagen fibrils rather than collagen itself. As a result, accurate determination of fibril width may be more challenging in contexts where proteoglycan abundance is reduced or elevated. Additionally, very small fibrils (e.g., <50 nm in diameter) may fall below the resolution achievable with the current 4-5× expansion factor. Taken together, these considerations highlight opportunities for further optimization of collagen fibril labeling and detection. For example, we found that increasing oxidation times improved collagen fibril labeling in bovine Achilles tendon sections. Moreover, resolving smaller fibrils may benefit from the use of higher expansion factor gels such as TREx (*58*) or MAGNIFY (*31*), in combination with imaging modalities offering enhanced resolution such as SIM-ExM (*59*). Overall, we anticipate that this approach, as implemented here or with further optimization of labeling and imaging conditions, will be universally applicable to the analysis of tendon architecture across developmental stages and disease contexts.

Elastic fibers have been proposed as key components of the tendon fascicular matrix, contributing to tendon mechanical properties (*16*), and potentially mediating the transmission of mechanical signals from collagen fibers to adjacent tenocytes (*60*). Support for this hypothesis has been based on the proximity of elastic fibers to tenocytes, and on the fact that perturbation of these fibers alters the bulk tendon mechanical properties. In our study, we identify a distinct elastic fiber network within collagen fiber bundles, in which fibers form interconnected networks and interface with sheet-like tenocyte regions. Interestingly, we observed a subset of fibers exhibiting a tube-like morphology, consistent with potential microfibrils surrounding an elastin core, which spanned the cellular regions of the tendon, either traversing cells or being closely wrapped by a neighboring cell. These observations raise the possibility that elastic fibers comprise two structurally and potentially functionally distinct populations: a more abundant thicker population that contributes to the structural organization of interconnected cellular regions, and a second population that spans fiber bundles, possibly facilitating mechanical signal transmission to nearby cells. Studying the formation of these fibers at larger length scales throughout tendon development may help elucidate the function of these different elastic fiber populations.

In our previous work, we demonstrated that loss-of-tension causes increased expression of catabolic gene programs, which leads to ECM remodeling and a weakening of the tissue (*40*). These changes were linked to epigenetic regulatory mechanisms, where a broad loss of chromatin accessibility was observed after de-tensioning, and overexpression of Yap/Taz *in vitro* helped prevent activation of the catabolic gene program while maintaining chromatin accessibility of Yap/Taz binding genes even after de-tensioning. In this study, we demonstrate that the loss of mechanical tension is associated with changes in carbohydrate-rich volume throughout the tendon, accompanied by an overall increase in glycoprotein content, which may be related to the changes in catabolic gene programs. Consistent with these results, previous studies have reported elevated levels of proteoglycans and GAGS in tendinopathic tissues (*61*). Despite these compositional changes, we did not observe a significant alteration in the number of branches, and thus in the number of collagen fiber bundles. We therefore propose that the observed increase in carbohydrate levels might represent greater cell-matrix adhesion as cells respond to the loss of the mechanical cue, given the variety of glycoproteins involved in focal adhesion maturation and establishment, such as heparin and chondroitin sulfate (*62*). Immunolabeling of focal adhesion components or proteoglycans associated with cell adhesion would shed more light into the molecular composition of these FLARE identified regions. Since FLARE-ExM is compatible with immunolabeling, these studies would represent an interesting future direction.

High resolution images of ruptured human Achilles tendon tissue remain rare in the literature and existing studies have largely focused on collagen fiber organization while providing limited insight into cellular architecture (*63*). Here, we present an initial examination of both collagen and cellular structures in ruptured human Achilles tendons. We observed an increase in cellularity across the tissue along with increased disorder in the collagen fiber regions and a loss of cell-cell and cell-matrix connections. Cells within these disrupted regions exhibited nuclei that lacked the typical elongated morphology of tenocytes, and instead displayed irregular nuclear shapes, a phenotype commonly associated with disruption of the nuclear lamina or reduced tension (*64*, *65*). Such changes are consistent with the altered mechanical environment resulting from extracellular matrix disruption. In addition, chromatin was enriched at the nuclear periphery, mirroring our previous observations from super-resolution imaging of tenocytes isolated from tendinopathy patients (*57*). Together, these findings indicate that FLARE-ExM can recapitulate key cellular features previously observed in isolated cells while enabling their visualization within intact tissue, thereby revealing coordinated changes in both cellular and extracellular matrix architecture.

The optimized protocol developed here for dense and mechanically stiff tendon tissue is likely to be relevant as researchers continue to adapt ExM to other dense and structurally complex tissue types, such as cartilage and bone. In this study, we were able to observe the ultrastructure of adjacent tissues, including muscle, which remained connected to the tendon explants. In this work, we expanded whole tendon explants from >10-week-old mice by carefully controlling the expansion rate and incorporating enzymatic digestion steps when necessary. Of note, we incorporated Elastase digestion, given the presence of elastic fibers within tendon tissue, and proteoglycan digestion, which improved the passivation of the ruptured human Achilles tendon sample. These enzymes, which are less commonly used for ExM compared to protease K and collagenase, highlight the principle of applying a targeted approach for improving tissue expansion. It is possible that the use of active diffusion strategies, such as with stochastic electric transport (*66*), or longer incubation times at each serial dilution step may allow more uniform expansion of tendons and other tissue types without the use of additional enzymatic digestion.

Overall, we demonstrate a robust approach for resolving fine cellular and extracellular architecture across tendon tissue, allowing for the interrogation of tendon cell biology questions across multiple spatial scales. We expect FLARE-ExM will provide a useful framework for linking tendon ultrastructure to mechano-epigenetic regulation, opening new avenues for understanding, and targeting, the structural and epigenetic changes that underlie tendinopathies.

## Materials and Methods

### Animals

All animal procedures were approved by the institutional Animal Care and Use Committee. The following mice were used: CD1 C57BL/6 with LifeAct-GFP for most samples, with the mouse tail tendon explants obtained from a CD1 C57BL/6 Myh9^HET^ and Myh10^KO^ mouse. The Bovine Achilles tendon was obtained from a 3 month old cow, which were frozen in OCT and cryo-sectioned into 50 µm sections, stored in 1×PBS at 4°C. Human ruptured Achilles tendon from a 46 year old male were obtained from the Penn Achilles Tendinopathy Center for Research Translation (PAT-CORT), which were frozen in OCT, cryo-sectioned into 100 µm sections, then fixed for 15 minutes in 4% paraformaldehyde (PFA) in PBS. Ruptured sections were washed three times in PBS, then stored in PBS at 4°C.

### FDL Resection Surgeries

We performed FDL tendon resection using our previously established protocol (*40*), described briefly as follows: Sterilely prepped mice (3 month old) were anesthetized and given pre-surgical analgesia. After exposing the distal FDL, a 3mm segment proximal to the bifurcation point of the tendon was resected to cause de-tensioning of the proximal region of the tendon nearest to the tibia. The skin was closed with 4-0 nylon sutures, and the animal was returned to unrestricted cage activity. Mice were then euthanized 1- or 6-days post-surgery, and both the surgery limb and the non-resected (control) hindlimbs were harvested and fixed in formalin for 4 days at 4°C. Limbs were then placed in 1×PBS buffer and stored at 4°C until use.

### Expansion Microscopy

Mouse tendons, excised from formalin-fixed hindlimbs, and other tendon cryosections were placed in a monomer incubation solution (5-10% (w/v) Sodium Acrylate, 20% (w/v) Acrylamide, 0.05% (w/v) Bis-acrylamide, 4% (w/v) paraformaldehyde, 1×PBS) supplemented with 0.1% (v/v) Triton-x 100 for all samples except the Bovine Achilles tendon cryosections. The samples were incubated in the monomer solution for 1-3 days on a gentle nutator at 4°C. Tendons were then placed on a glass slide with a brush or metal spatula, and a 22×22 mm coverslip was placed on top of each group of tendons. A gel mix was made by supplementing monomer solution, with or without 4% PFA, with 0.1% (w/w) VA-44 solution. The gel mix was pipetted slowly between the coverslip and glass slide until the gel mix surrounded the tendon explants and covered the area between coverslip and glass slide. For the Bovine Achilles tendon explants, a set of two 22×22 mm No. 1.5 coverslips were placed as spacers beside the cryosection and between the top coverslip and the glass slide, whereas the rest of tendon samples had no spacers. The gelation assemblies were then placed in a sealable container that was backfilled with nitrogen gas for ∼1 min. Sealed containers were placed in a water bath set to 45°C, and samples were gelled for 2-3 hours. Excess gel was cut away with a sharp blade, and the remaining tendon-containing gel was measured with a ruler. Bovine Achilles tendon samples were placed in 1mL of denaturation solution A (200 mM Sodium Dodecyl Sulfate, 200 mM Sodium Chloride, and 50 mM Tris (pH 9.0)), and the tendon explants, along with the human ruptured Achilles tendon sections, where placed in denaturation solution B (200 mM Sodium Dodecyl Sulfate, 400 mM Sodium Chloride, 20 mM Calcium Chloride and 50 mM Tris (pH 9.0)). Samples were then denatured at 65-70°C for 2-3 days, then at 85-90°C for 1-3 days. For specific details, see Table S1.

A collagenase digestion step, which helped bring uniformity across thicker tendons but at the loss of ECM details, was performed on the gelled Achilles and FDL tendon samples in Fig. 3 and 4, where gelled samples were placed in collagenase VII solution (500 U/mL of Collagenase type VII, 500 mM Sodium Chloride, 40 mM Calcium Chloride and 100 mM Tris (pH 8.0)) and digested for 24 hours at 37°C with gentle nutating before denaturation. For specific details, see Table S1.

After the denaturation step, samples were washed three times with 20 mL of 100 mM Tris (pH 8), three times for 10 minutes with gentle nutating at RT. Samples digested with collagenase were further treated with an Elastase solution (2.5 U/mL Elastase, 0.1 mg/mL Soy Bean Trypsin Inhibitor, 100mM Tris (pH 8)) for 2.5 hrs at 37°C, with gentle nutating. Samples were then washed with 20 mL of high salt buffer (250 mM NaCl with 50 mM Tris (pH 8)), three times for 10 minutes and stored in the cold room, until processed for FLARE within a week. For specific details, see Table S1.

The Ruptured human Achilles tendon 100 µm cryosections were processed the same way as the digested mouse tendon explants, except that the post-denaturation digestion solution included 50 mM of Sodium Acetate and 0.1 U/mL of chondroitinase ABC. For specific details, see Table S1.

### FLARE-ExM on tendons

FLARE staining of expanded tendons was adapted from previous works (*32*, *67*), and summarized as follows: Tendons were washed twice with 10 mL of high salt acetate buffer ( 1M Sodium Chloride and 100 mM Sodium Acetate (pH 5)) for 10 minutes each time with gentle nutating at RT. Samples were then oxidized for 1 to 4 hours with 20 mM Sodium Periodate in high salt acetate buffer at 37°C with gentle shaking. Samples were then washed three times with 100 mM Sodium Acetate (pH 5), 10 minutes each time, with gentle nutating at RT. Samples were then incubated in 5-10 µg/mL of ATTO TEC 565 hydrazide in 100 mM sodium acetate (pH 5) for 2 hours, covered in foil with gentle nutating at RT. After 2 hours, samples were sealed, covered in foil, and incubated at 4°C overnight with gentle nutating. Samples were then reduced with the addition of 20 mM of Sodium cyanoborohydride to the existing solution, in a sealed container for 30 minutes with gentle nutating at RT. Samples were then washed twice with 100 mM sodium acetate (pH 5) for 10 minutes each time, and a third time for 1 hour. Samples were then washed with PBS for 10 minutes with gentle nutating at RT. Sample were then placed in 6-12 µg/mL of ATTO TEC 647N NHS ester for 2 hours with gentle nutating. Samples were then sealed, covered in foil, and incubated overnight with gentle nutating at 4°C. Samples were then washed twice with 100 mM Tris pH 8, 10 minutes each time with gentle nutating at RT. Samples were then incubated in 1:1000 SybrSf in 100mM Tris (pH 8), either for 1-4 hours with gentle nutating at RT, or overnight at 4°C with gentle nutating. Bovine Achilles tendon sections were expanded in water, replacing the water three times every 20 minutes, with gentle nutating at RT during each step. For mouse tendon explants, samples were incubated in 20 mM Tris (pH 8), then in 4mM Tris (pH 8), then twice in DI waters, each time for 20-30 min at RT and with gentle nutating. For specific details, see Table S1.

### Immunofluorescence on expanded tendons

For immunolabeling, expanded and FLARE-stained carbohydrate samples of bovine Achilles tendon cryosections were first incubated in 1% Triton-x 100 in 1×PBS (PBST) for 1 hour at RT. Samples were then incubated in 10 µg/mL of Mouse anti-Elastin in PBST for 24 hrs at 37°C. Samples were washed three times with PBST, 1 hr each time at 37°C. Samples were then incubated in 10 µg/mL of Donkey anti-Mouse conjugated to Biotin in PBST for 24 hrs at 37°C. Samples was washed three times with 1×PBS, 1 hr each time at 37°C. Samples were then incubated in 10 µg/mL conjugated to streptavidin in PBST for overnight at 4°C. Samples were incubated in excess water for 20 minutes at RT with gentle shaking, then in an imaging buffer (2% Methanol, 1mM Methyl Viologen, 1mM L-ascorbic acid, 30 mM Tris pH 8) for 20 minutes at RT with gentle shaking. Sample was then image by confocal.

### Nuclear staining of unexpanded and expanded mouse tail tendon explants

An unexpanded mouse tail tendon was placed in incubation monomer with 1 µg/mL of Hoechst 33258 overnight at 4°C with gentle shaking. After following the steps for gelation as described above, the sample was then imaged by confocal, and the rest of the expansion protocol was followed.

After denaturation and washing of the tendon with 100 mM Tris (pH 8) as described above, the tendon was then incubated in 10 µg/mL of Hoechst 33258 in 1×PBS for two hours at RT. The sample was then expanded in 20mM Tris (pH 8), 4 mM Tris (pH 8) and water as described above. The nuclei of the tendon were imaged on a confocal, and the aspect ratios of the nuclei were compared to that of the unexpanded nuclei. For specific details, see Table S1.

### FLARE on explanted patellar tendon for correlative imaging

To obtain a dense label of the tendon tissue structure for pre- and post-imaging of the patellar tendon, without extensive perturbation of protein anchoring to the gel, we permeabilized the tendons for 1 hour in 0.5% Triton-x 100 in 1×PBS, then incubated the tendon in 0.1-0.01 µg/mL of ATTO647N NHS ester in the same solution for another hour. Samples were then washed three times with PBS, 10 minutes each time. The patellar tendons were then incubated in monomer solution and gelled in the same manner as the rest of the mouse tendon explant. Before collagenase digestion or denaturation, the protein stain for the whole tendon was imaged at a low magnification, with specific regions of the tendon imaged at a higher magnification. The remainder of the expansion protocol was then followed. The expanded patellar tendons were first incubated in PBS for 20-30 minutes, three times, before following the FLARE procedure for protein staining post-expansion. The tendons were then imaged again on the confocal, and correlative images were compared to each other. For specific details, see Table S1.

### Confocal microscopy

All microscopy images were obtained on a ZEISS Laser Scanning Confocal Microscope (LSM980) using either a 5× 0.4 NA air objective lens, a 20× 0.8 NA air objective lens, or a 40× 1.1 NA water immersion lens. The images in Fig. 1B. for pre and post expansion were obtained using a Pixel 6a camera phone. A linear Wiener filter deconvolution was applied to the higher magnification images using the LSM plus feature for all images except for those in Fig. 2B, Fig. 3B (left) and Fig. S8.

### Full Width Half Max (FWHM) measurements of expanded collagen fibrils

To measure the FWHM of the bovine Achilles tendon fibrils that appeared in the carbohydrate stain horizontal view, we used a custom MATLAB script, first generated with Microsoft AI copilot (GPT-5 model) then edited to incorporate additional processing steps, briefly described as follows: The image was smoothed with a gaussian filter, then binarized and divided using watershed. Small (4 pixel) and large (80 pixel) objects were filtered out, and the centroid was obtained for the remaining puncta. The raw intensity values within a set box around each centroid were fit to a 2D gaussian curve, and any results that provide centroids outside of the specified Region of Interest (ROI) for fitting were removed. The x and y sigma’s from the fit result were multiplied by 2.3548 and the pre-expansion correct pixel size to obtain the FWHM of each puncta.

To measure the FWHM of the detected fibrils within sagittal fields of view for each tendon sample, we searched for areas within a field of view that presented distinguishable and linear fibrils over a length of ∼1.5 microns. Using Fiji, we obtained a line plot measured across the individual fibril, with carbohydrate intensity signal averaged across the ∼1.5-micron distance (100 pixels). The line plot values were fitted to a gaussian curve and, If the R^2^ value for the curve was greater than 0.9, the sigma value was recorded and multiplied by 2.3548 to measure the FWHM of the fibril. Values were then corrected to the expansion factor of each sample. A total of 10 fibrils were measured per field of view (FOV), 1 FOV per tendon in Fig. S4B and 3 FOV per tendon per sample in Fig. 4C.

### FLARE carbohydrate enrichment in resected vs. uninjured FDL tendons

To determine the amount of carbohydrate signal enrichment for the FDL resection model in Fig. 4D, raw confocal images of a 250×250×20 µm³ tendon volume was maximum intensity projected for all FLARE channels (Fig. S7&8). The average carbohydrate signal intensity over the projected area was subtracted by the average intensity over an area of the same gel that contained no tendon. Each tendon value was then divided by the average value of the uninjured control conditions of the same day post-surgery.

### Ilastik segmentation of tendon structures

Tendon cells and fiber segmentation was performed using the ilastik pixel classification workflow (*54*), where sparse strokes were used to indicate each class. For the bovine Achilles tendon fibrils (Fig. 2C), strokes of the FLARE-stained carbohydrate channel indicated fibril or background. For the mouse Achilles tendon explant 3D stack (Fig. 3C&D, Fig. S6D and Movie S1), strokes on FLARE-stained protein, carbohydrate and nuclear channels indicated: nuclei, protein-rich region, carbohydrate-rich region, collagen fiber-bundle region, and elastic fibers region. For The FDL tendon resection model horizontal 2D measurement (Fig. 4E&F and S10A&B), a 100×100 µm² region of interest, which excluded the more brightly stained paratenon region, for each pair of FDL tendons (i.e., uninjured and resected) were processed together, where strokes on FLARE-stained protein, carbohydrate and nuclear channels indicated: nuclei, protein-rich region, carbohydrate-rich region, collagen fiber-bundle region, and no tendon region. For the FDL resection model 3D horizontal stacks (Fig. 4G-I, and Movie S2), strokes on FLARE-stained protein and carbohydrate channels indicated: nuclear region, protein-rich region, carbohydrate-rich region, and collagen fiber region.

### MATLAB processing and analysis of segmented regions

For nuclear aspect ratio measurements (Fig. S1C), nuclei were segmented from maximum intensity projected nuclear stain of the same mouse tail tendon explant imaged before and after expansion. Segmentation was done using the Segment anything model (SAM) region boundaries brush in MATLAB image segmenter. Nuclei that were distinct from other nuclei and away from the image border were selected for the analysis.

For ilastik segmented mouse Achilles tendon explant regions (Fig. 3C&D, Fig. S6D and Movie S1), small objects (less than 2500 voxels) were removed before rendering of volumes. For ilastik segmented FDL tendon resection model 3D horizontal regions (Fig. 4G-I, and Movie S2), each region was subjected to a morphological closing, then opening, using a 3-voxel radius sphere, followed by removal of small objects (<2500 pixels). The protein and carbohydrate-rich region fraction was calculated by dividing the total volume of both regions per sample, over the whole tendon volume in each field of view. To determine the number of cell sheets within each sampled tendon volume, the raw carbohydrate channel was superimposed with the segmented regions to help visualize dimmer connections that may not have been completely segmented by ilastik. Sheets were then manually counted per tendon image stack.

For ilastik segmented FDL tendon resection model 2D horizontal regions (Fig. 4E&F and S10A&B), nuclear, carbohydrate-rich and collagen-bundle regions were subject to morphological closing, then opening, using a 3-pixel radius disc, followed by removal of small objects (<2000 pixels). The same procedure was done for the protein-rich region and no tendon region, except a 20-pixel radius disk was used instead. All segmented nuclei at the ROI border were cleared. The area and circularity of the remaining nuclei were calculated. Protein and carbohydrate-rich region fractions were calculated by dividing the total area of both regions per sample, over the whole tendon region in each field of view. Branchpoints for cell connections originating at the cytoplasm surrounding each nucleus was determined by first skeletonizing a dilated combined region of the carbohydrate and protein-rich areas (dilated with a 20-pixel disk), removing branches with a length less than 50 pixels, then counting the number of branchpoints that fell within 200 pixels of each nucleus (i.e., branchpoints within a region generated by dilating the nuclear region with a disk with a 200 pixel radius).

3D model representation in Movie S1 and S2 were generated using a custom MATLAB script, then stitched together using a script generated by Microsoft AI copilot (GPT-5 model).

### Statistical Analysis

For determining significance between different collagen fibrils of different tendon types in Fig. S4A, a multivariate ANOVA was applied between all tendon samples, with n = 10 fibrils per sample. For the uninjured and resected FDL tendon measurements, obtained from the left (resected) and right (uninjured) hindlimb FDL tendon explants for 3 mice per time point (i.e., 1 day or 6 days post-surgery), significance was determined by a pairwise 2-way t-test between the mean-values of uninjured and resected tendons. All statistical tests were applied using MATLAB. A result was considered significant for p-values less than 0.05.

## Acknowledgments

The authors acknowledge Dr. Andrea Stout and the Cell and Developmental Biology (CDB) Microscopy Core Facility in the Perelman School of Medicine at the University of Pennsylvania for access and training for confocal imaging. The authors acknowledge Dr. Casey Humbyrd, Dr. Lou Soslowsky and the Penn Achilles Tendinopathy Center of Research Translation (PAT-CORT) at the University of Pennsylvania for providing human ruptured Achilles tendon tissue.

## Funding

National Institutes of Health grant T32AR053461 (MW)

National Institutes of Health grant P50AR080581 (RM, SCH, ND, ML)

National Institutes of Health grant R01AR079224 (SCH, ML)

National Institutes of Health grant R01GM133842 (ML)

National Institutes of Health grant RM1GM136511 (ML)

National Institutes of Health grant R35GM152111 (ML)

National Science Foundation grant CMMI1548571 (ML)

## Author contributions

Conceptualization: MAW, ML, ND, RM

Methodology: MAW, TM, XJ, ZL, YH, MKE

Investigation: MAW, TM, XJ

Visualization: MAW

Supervision: ML, RM, ND, SCH

Writing of original draft: MAW, ML

Review & editing: MAW, ML, ND, RM, TM, XJ, ZL, YH, MKE, SCH

## Data and materials availability

Imaging data is available upon request, as it is too large to deposit in a data repository. All other data values used in making the plots in the main and supplementary figures have been deposited to Figshare (https://doi.org/10.0.23.196/m9.figshare.31980525).

## Competing interests

Authors declare that they have no competing interests.

## Notes

### Competing Interest Statement

The authors have declared no competing interest.

