## Supplementary Information for "Expansion Microscopy reveals changes in cellular and extracellular structures between healthy and perturbed tendons"

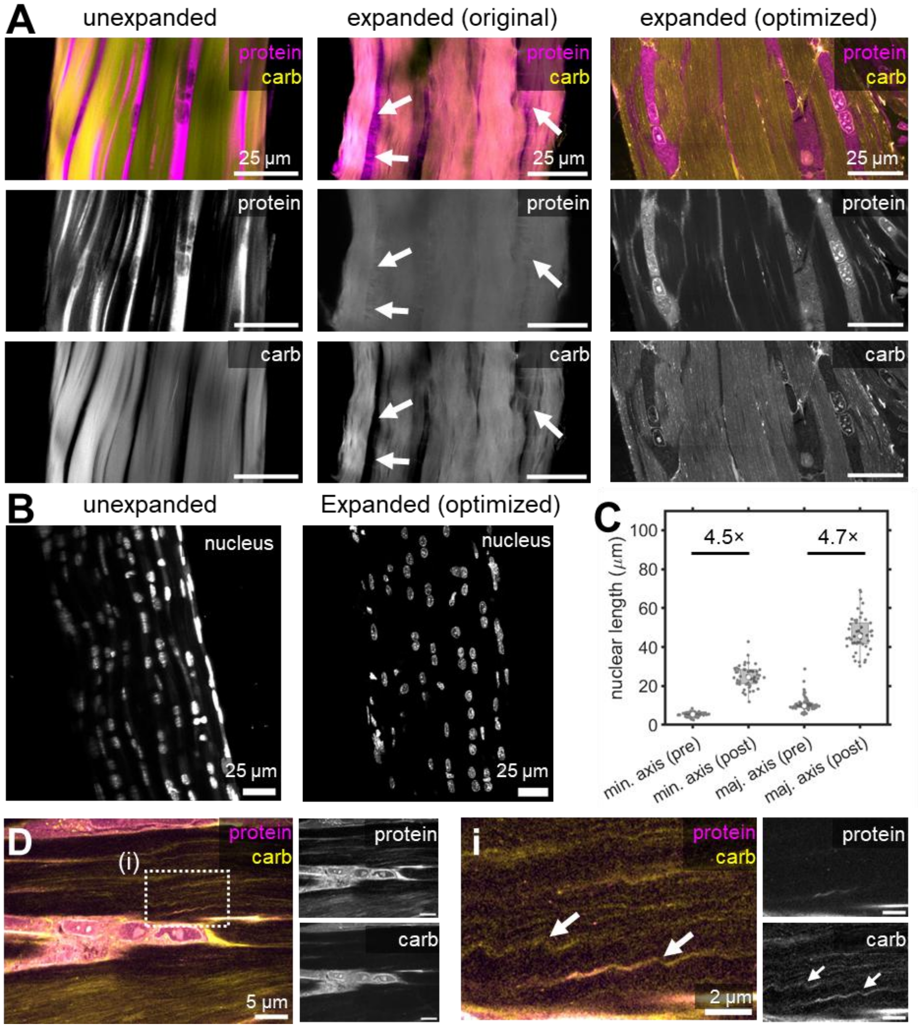


**Fig. S1. Adapting Expansion Microscopy for the visualization of Mouse tendon explants. A.** FLARE staining of proteins (magenta) and carbohydrates (carb, yellow) of unexpanded (left), attempted expansion with the original MAP protocol (middle), and expanded mouse tail tendon explants using the optimized protocol (right). Arrows indicate cells in the original MAP protocol that appear distorted and non-uniformly expanded. **B.** Comparison of Hoechst labeled nuclei diameters of the same mouse tail tendon explant, before and after expansion with the optimized protocol. **C.** Lengths of the major and minor axes of nuclei of the mouse tail tendon explant in **(B.)**, along with the ratio between the pre and post expansion median measurement. **D.** FLARE staining of proteins (magenta) and carbohydrates (yellow) of expanded mouse Flexor Digitorum Longus tendon explant cells, which show "crinkling" in certain regions near the cells (indicated by white arrows in zoomed in view in **i.**). Image in **(B, left)** was rotated by 45° using bilinear interpolation to align with axis of the image of **(B, right)**. Images in **(A.)** and **(D.)** were subject to a 2-pixel radius median filter.

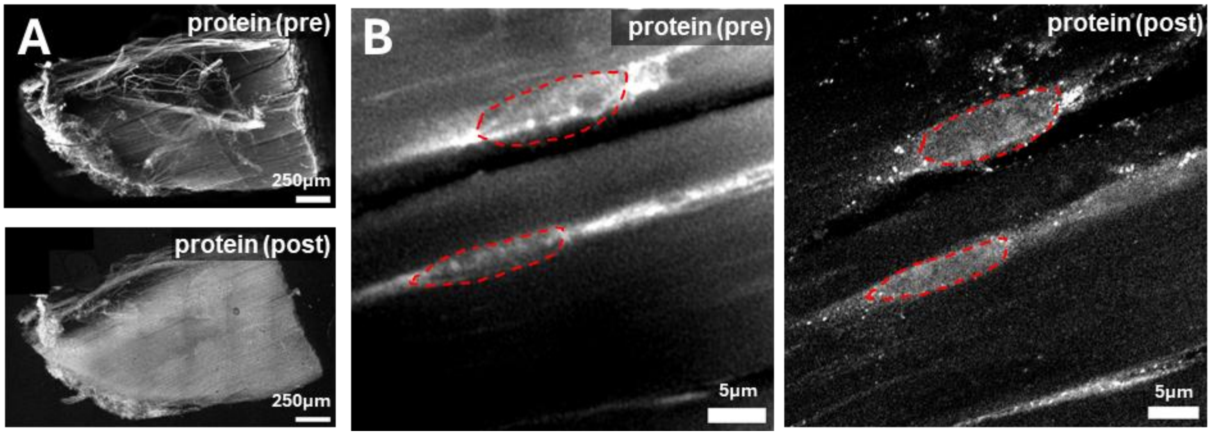


**Fig. S2. Correlative imaging of expanded mouse patellar tendon explants treated with collagenase before denaturation and elastase after denaturation. A.** Correlative imaging of a whole mouse patellar tendon explant FLARE-stained for proteins before expansion (top) and after expansion (bottom), maximum intensity projected over 58µm. **B.** High-magnification image, maximum intensity projected over 1.25µm, of cells within the pre- and post-expanded patellar tendon explant in **(A.),** with a red outline indicating the same traced cell nuclei perimeter between pre- and post- expansion. Pre-expansion images in **(A.)** and **(B.)** were rotated 35° using a bilinear interpolation to achieve an approximate alignment with post-expansion images.

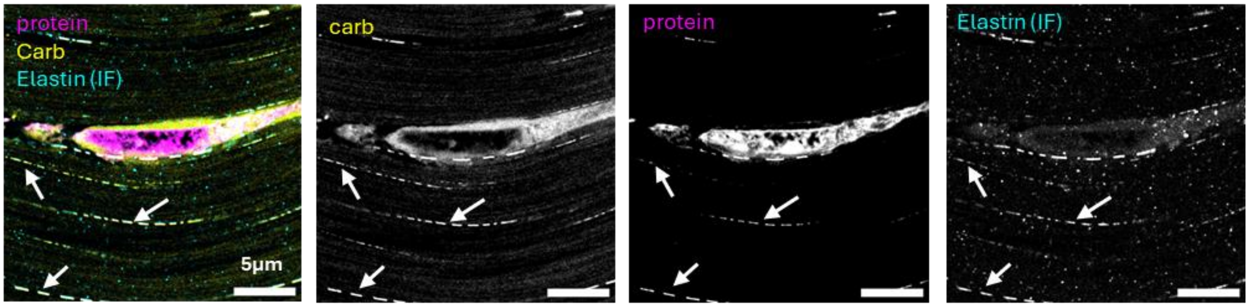


**Fig. S3. Immunofluorescence of expanded tendons.** IF staining of Elastin (cyan) after FLARE staining of proteins (magenta) and carbohydrates (carb, yellow) in an expanded bovine Achilles tendon 50 µm cryosection.

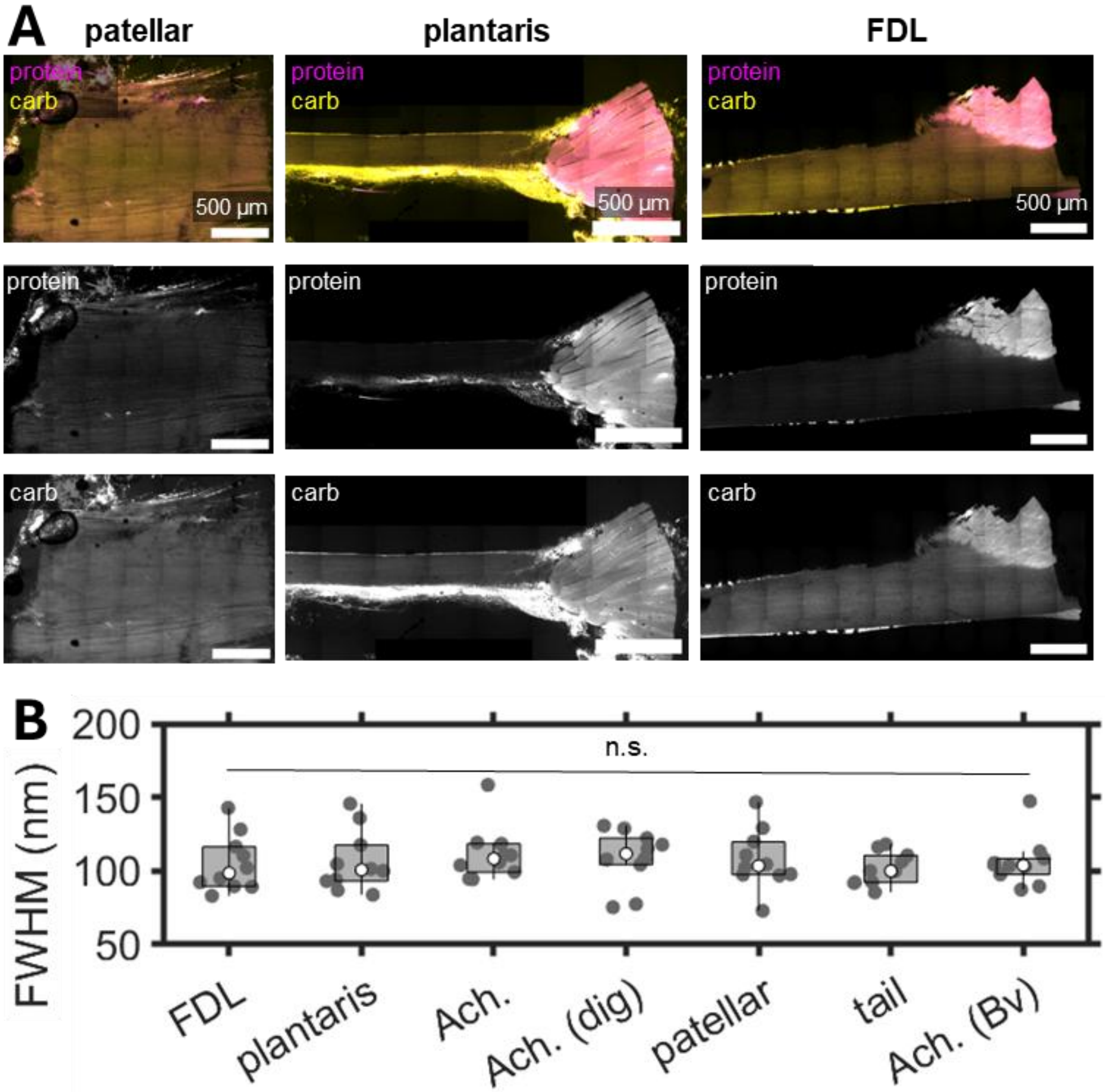


**Fig. S4. Example images of FLARE-ExM on whole tendon explants. A**. Example images of FLARE-stained proteins (magenta) and carbohydrates (yellow) for expanded mouse patellar (left), plantaris (middle), and *Flexor Digitorum Longus* (FDL, right) mouse tendon explants**. B.** box plots for the FWHM measurements of FLARE-stained carbohydrate fibril-like structures in a sagittal view of the FDL, plantaris, Achilles (Ach.), digested (dig.) Achilles, patellar, tail tendon explants, along with measurements from the bovine (Bv) Achilles tendon 50 µm cryosection sagittal view in **Fig. 2A.** Images in **(A.)** were subject to a 2-pixel radius gaussian filter.

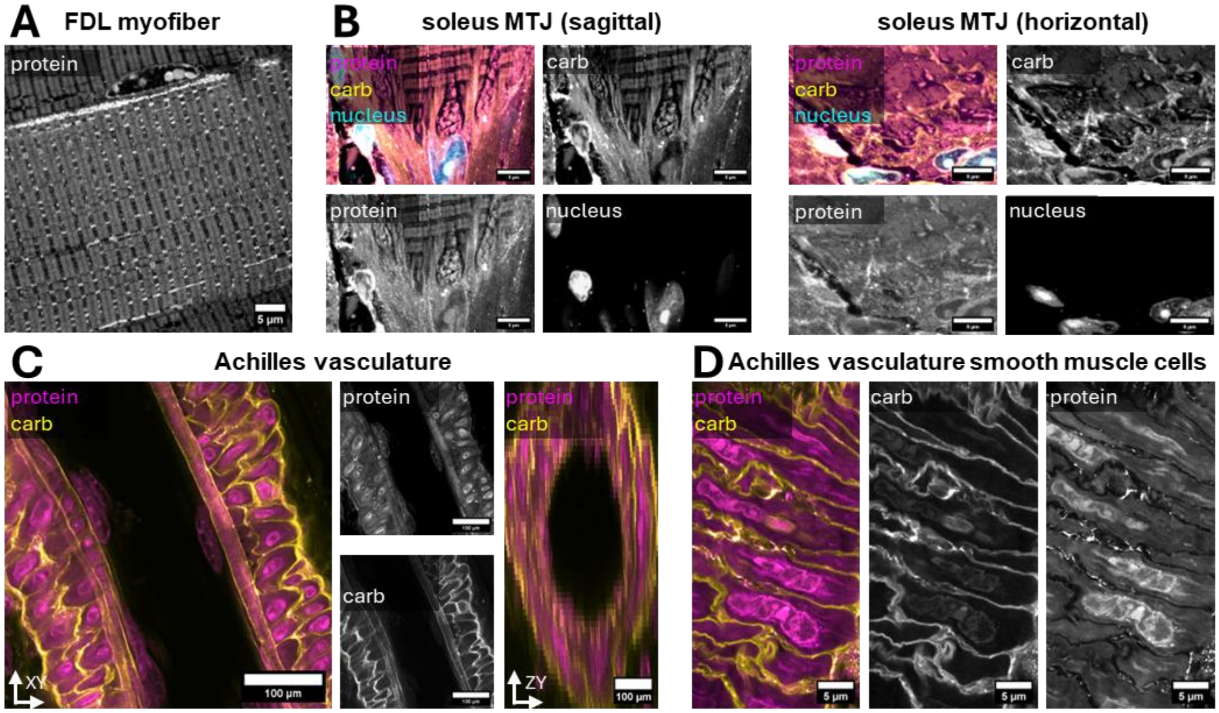


**Fig. S5. Visualizing adjacent tissues to expanded mouse tendon explants. A.** FLARE staining of proteins of the *Flexor Digitorum Longus* (FDL) muscle, showing organelle-like structures the myofiber sarcomeres. **B.** FLARE staining of protein (magenta), carbohydrates (carb, yellow) and of nuclei (cyan) of the expanded myotendinous junction (MTJ) between the Soleus and the Achilles tendon, either in a sagittal view (left) or horizontal view (right). **C.** FLARE staining of proteins (magenta) or carbohydrates (carb, yellow) of the expansion of a major artery found near the Achilles tendon seen as a longitudinal section in the XY imaging plane (left), or a cross-section in the ZY plane (right). **D.** High-magnification of FLARE-stained proteins (magenta) or carbohydrates (carb, yellow) of the vascular smooth muscle cells of the major artery seen in **(C.)**.

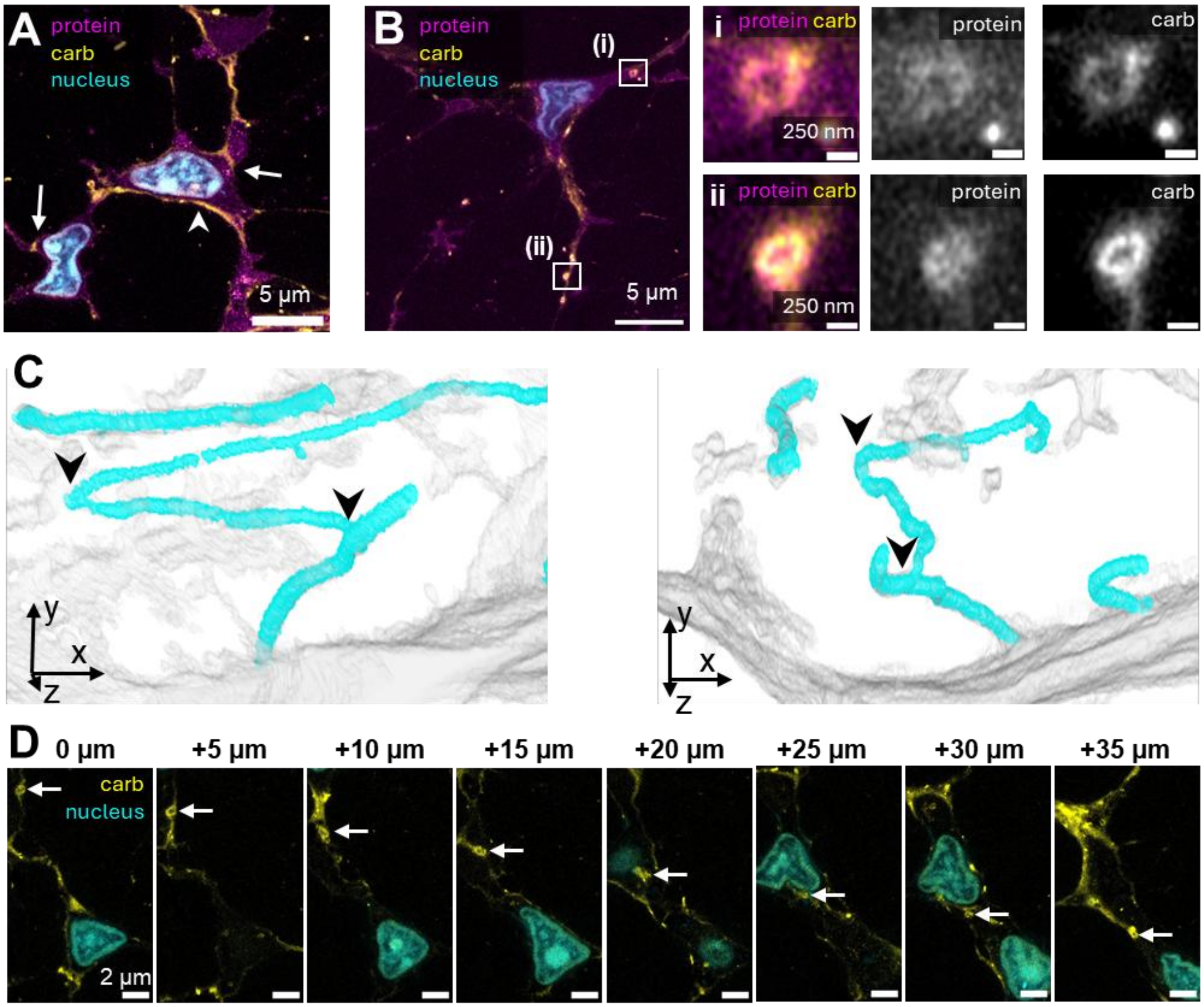


**Fig. S6. FLARE-ExM of mouse Achilles tendon explants reveals carbohydrate rich structures between cells. A.** FLARE staining of carbohydrates (carb, yellow), protein (magenta) and nuclei (cyan) of the Mouse Achilles tendon explant in **Fig. 3A,** with arrows indicating carbohydrate-rich regions between cells and the arrowhead indicating a carbohydrate-rich region between a cell and an adjacent collagen fiber bundle. **B.** FLARE staining of carbohydrates (carb, yellow), protein (magenta) and nuclei (cyan) of the Mouse Achilles tendon explant in **Fig. 3A**, with zoomed in views **i.** and **ii.** showing ring-like structures within the cellular region of the tendon. **C.** Zoomed in view of segmented elastic fibers (cyan) in the context of all non-collagenous segmented regions (gray) from **Fig. 3C** with elastic fiber branching points indicated by arrowheads. **D.** Same field of view of the mouse Achilles tendon cellular region in **Fig. 3B** imaged at different depths, with the arrow indicating in each panel the progression of a ring-like structure either in contact or passing through multiple cell regions.

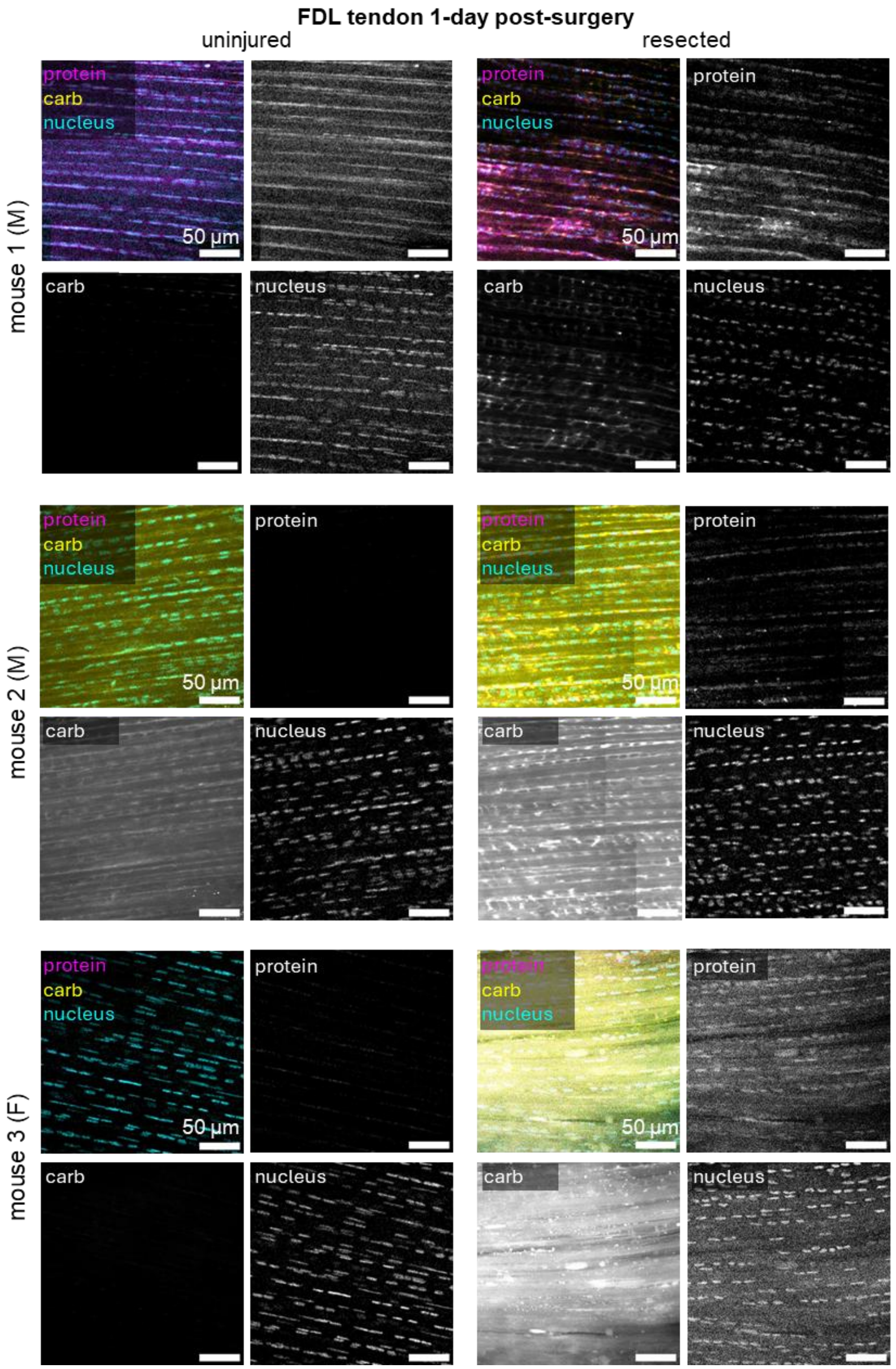


**Fig. S7. Sagittal views of FLARE-ExM Flexor Digitorum Longus (FDL) tendon explants 1-Day post-surgery.** Maximum intensity projection of a 20 µm z-depth of FLARE-stained protein (magenta), carbohydrate (yellow) and nuclei (cyan) for the uninjured (left) and resected (right) FDL tendon explants for three mice sacrificed 1-day post-surgery. M indicates Male and F indicates Female. All images were subject to a 1-pixel radius gaussian filter and are displayed with the same contrast levels.

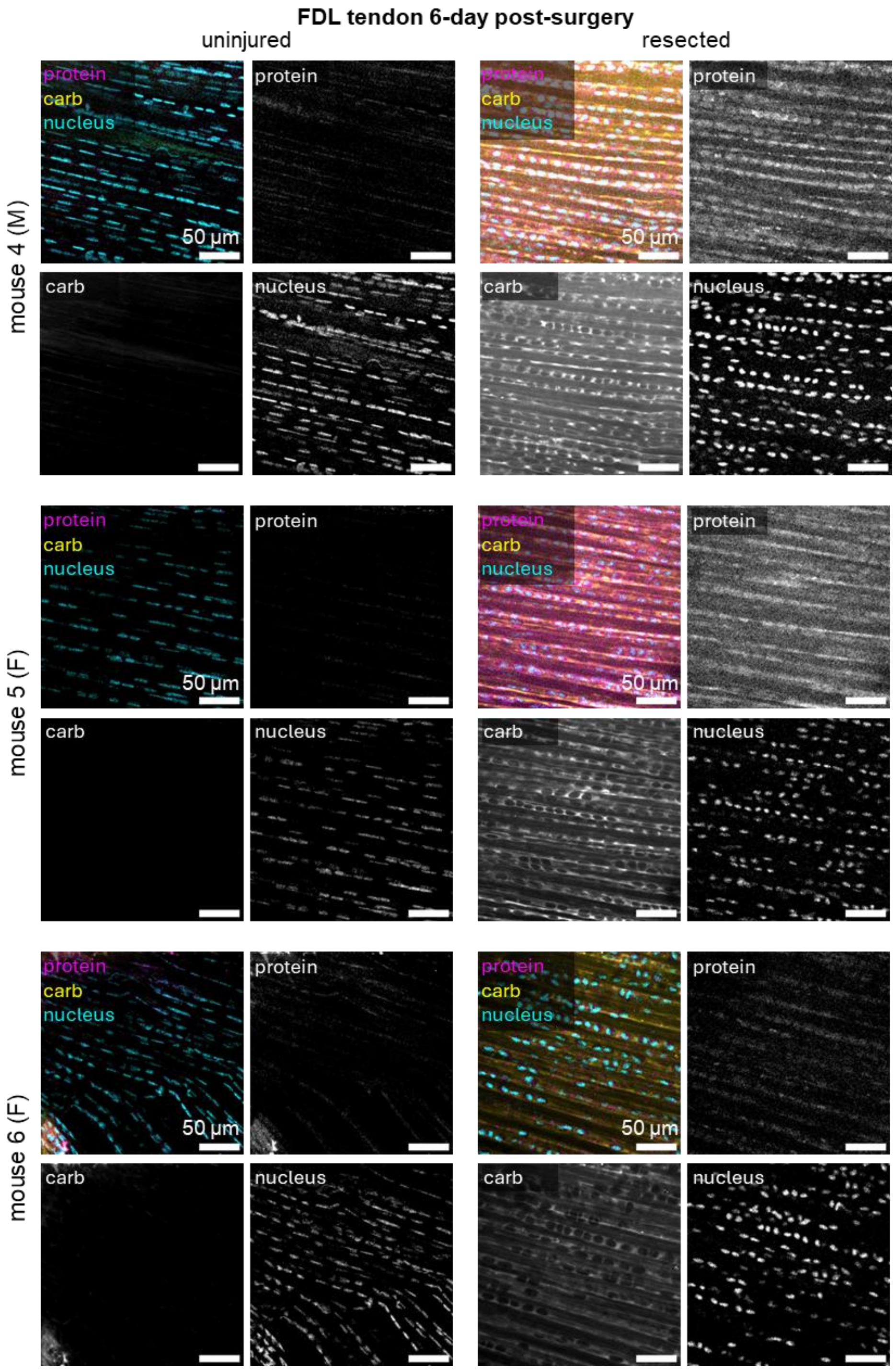


**Fig. S8. Sagittal views of FLARE-ExM Flexor Digitorum Longus (FDL) tendon explants 6-Days post-surgery.** Maximum intensity projection of a 20 µm z-depth of FLARE-stained protein (magenta), carbohydrate (yellow) and nuclei (cyan) for the uninjured (left) and resected (right) FDL tendon explants for three mice sacrificed 6-days post-surgery. M indicates Male and F indicates Female. All images were subject to a 1-pixel radius gaussian filter and are displayed with the same contrast levels.

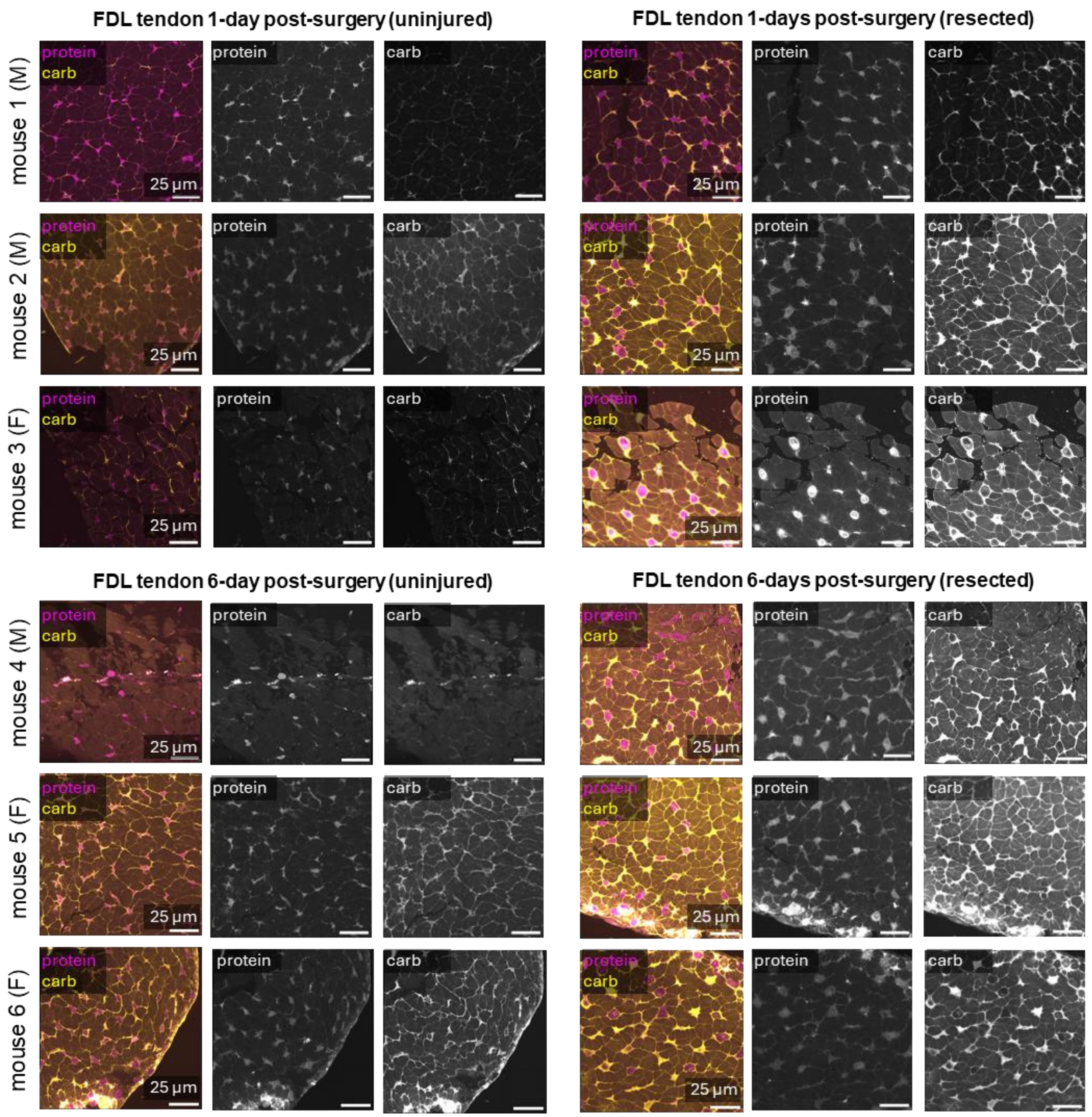


**Fig. S9. Horizontal views of FLARE-ExM Flexor Digitorum Longus (FDL) tendon explants 1- or 6-Days post-surgery.** FLARE stained protein (magenta) and carbohydrate (carb, yellow) for the uninjured (left) and resected (right) FDL tendon explants for three mice sacrificed 1-Day post-surgery in **A.** and 3 mice sacrificed 6-Days post-surgery in **B.** M indicates Male and F indicated Female. All images were subject to a 2-pixel radius gaussian filter and are displayed with the same contrast levels.

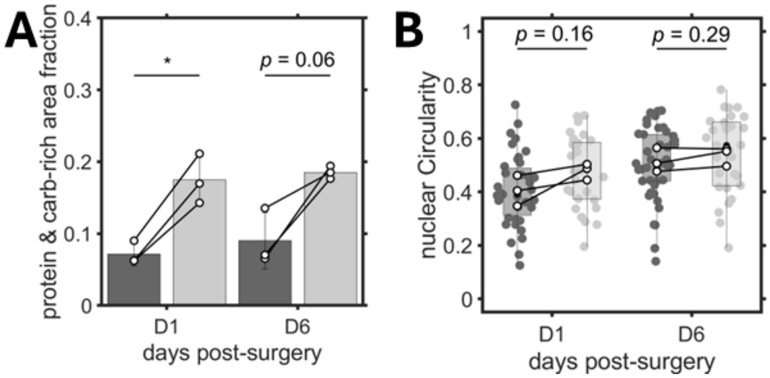


**Fig. S10. De-tensioned *flexor digitorum longus* (FDL) tendons shows higher protein- and carbohydrate-rich regions compared to uninjured tendons. A.** Bar plots for the fraction of protein- and carbohydrate (carb)-rich ilastik segmented horizontal view tendon regions of a 100×100 $\mu m^2$ Region of Interest (ROI) that excludes the paratenon region per tendon seen in **Fig. S9. B.** Box plot for circularity of ilastik segmented nuclei within a 100×100 $\mu m^2$ Region of Interest (ROI) that excludes the paratenon region per tendon seen in **Fig. S9.**

**Table S1.**

Expansion and FLARE labeling conditions per tendon sample.

| Fig, | Sp. | tnd. | type | Inc. mon. Acry. con. (%) | Inc. mon. TX-100 con. (%) | inc. mon. time (d) | 4% PFA in gel mix? | Coll., Ela.? | Den. Sol. | 70°C den. Time (d) | 90°C den. Time (d) | cABC? | Ox. Time (hr) | Carb stain con. (µg/ mL) | Prtn stain con. (µg/ mL) | Nuclear stain | Ex. F. |
|---|---|---|---|---|---|---|---|---|---|---|---|---|---|---|---|---|---|
| 1B S1D S4A | Ms. | FDL | explant | 10 | 0.1 | 3 | Yes | No | B | 3 | 3 | No | 2 | 10 | 10 | SybrSf | 4.8 |
| 1C | Ms | Pat. | explant | 5 | 0.1 | 3 | No | No | B | 3 | 3 | No | -- | -- | 10 | -- | 4.6 |
| 2A | Bv | Ach. | 50 µm section | 10 | 0 | 3 | Yes | No | A | 3 | 3 | No | 1 | 5 | 6 | SybrSf | 4.3 |
| 2B 2C | Bv | Ach. | 50 µm section | 10 | 0 | 2 | Yes | No | A | 3 | 3 | No | 4 | 10 | -- | SybrSf | 4.3 |
| 2D S1A (right) | Ms | Tail | explant | 10 | 0.1 | 1 | Yes | No | B | 2 | 1 | No | 2 | 5 | 10 | SybrSf | 4.3 |
| 2D S4A | Ms | Pat. | explant | 10 | 0.1 | 2 | Yes | No | B | 3 | 3 | No | 2 | 10 | 10 | -- | 4.7 |
| 2D S4A | Ms | Plant. | explant | 10 | 0.1 | 2 | Yes | No | B | 3 | 3 | No | 2 | 10 | 10 | -- | 4.9 |
| 2D S5 | Ms | Ach. | explant | 10 | 0.1 | 2 | Yes | No | B | 3 | 3 | No | 2 | 10 | 10 | -- | 4.3 |
| 3 S6 | Ms | Ach. | explant | 10 | 0.1 | 3 | Yes | Yes | B | 3 | 3 | No | 2 | 10 | 10 | SybrSf | 4.7 |
| 4 S7-9 | Ms | FDL | explant | 10 | 0.1 | 3 | Yes | Yes | B | 3 | 3 | No | 2 | 10 | 10 | SybrSf | 4.4-4.8 |
| 5 | Hu | Ach. | 100 µm section | 5 | 0.5 | 3 | No | Yes | B | 3 | 3 | Yes | 2 | 10 | 10 | -- | 5.0 |
| S1A (mid.) | Ms | Tail | Tendon | 10 | 0 | 1 | Yes | No | A | 3 | 1 | No | 2 | 2 | 10 | -- | -- |
| S1B | Ms | Tail | Tendon | 10 | 0 | 1 | Yes | No | B | 2 | 1 | No | -- | -- | -- | Hoechst | 4.6 |
| S2 | Ms | Pat. | explant | 5 | 0.1 | 3 | No | Yes | B | 3 | 3 | No | -- | -- | 10 | -- | 4.8 |
| S3 | Bv | Ach. | 50 µm section | 10 | 0 | 2 | Yes | No | A | 3 | 3 | No | 2 | 5 | 10 | -- | 4.3 |

**Abbreviations:** Fig.: Figure, Sp.: species, tnd.: tendon type, Inc. mon.: incubation monomer, Acry.: Sodium Acrylate, con.: concentration, TX-100: Triton-x 100, PFA: Paraformaldehyde, Coll.: Collagenase type VII, Ela.: Elastase type II, Den. Sol.: Denaturation Solution, cABC: chondroitinase ABC, Ox.: Oxidation, Carb: Carbohydrate, Prtn: protein, Ex. F.: Expansion factor, Ms: Mouse, Bv: bovine, Hu: Human, FDL: Flexor Digitorum longus, Pat.: Patellar, Ach.: Achilles, Plant.: Plantaris

**Movie S1.**

**Rotating Volume of ilastik segmented regions of FLARE-stained mouse Achilles tendon explant.** The segmented regions correspond to nuclei (blue), protein-rich (magenta), carbohydrate-rich (yellow) and elastic fibers (cyan). The volumes were scaled down by half to enable rendering in MATLAB.

**Movie S2.**

**Rotating Volume of ilastik segmented regions of FLARE-stained FDL resection tendon model.** The segmented regions correspond to nuclei (blue), protein-rich (magenta), carbohydrate-rich (yellow) for uninjured (left) and resected (right) FDL tendons 6-days post-surgery. The volumes were scaled down by half to enable rendering in MATLAB.
